# Salt-bridge Dynamics in Intrinsically Disordered Proteins: A trade-off between electrostatic interactions and structural flexibility

**DOI:** 10.1101/220392

**Authors:** Sankar Basu, Parbati Biswas

**Author notes:** **E-mail:** SB, PB.

## Abstract

Intrinsically Disordered Proteins (IDPs) are enriched in charged and polar residues; and, therefore, electrostatic interactions play a predominant role in their dynamics. In order to remain multi-functional and exhibit their characteristic binding promiscuity, they need to retain considerable dynamic flexibility. At the same time, they also need to accommodate a large number of oppositely charged residues, which eventually lead to the formation of salt-bridges, imparting local rigidity. The formation of salt-bridges therefore oppose the desired dynamic flexibility. Hence, there appears to be a meticulous trade-off between the two mechanisms which the current study attempts to unravel. With this objective, we identify and analyze salt-bridges, both as isolated as well as composite ionic bond motifs, in the molecular dynamic trajectories of a set of appropriately chosen IDPs. Time evolved structural properties of these salt-bridges like persistence, associated secondary structural ′order-disorder′ transitions, correlated atomic movements, contribution in the overall electrostatic balance of the proteins have been studied in necessary detail. The results suggest that the key to maintain such a trade-off over time is the continuous formation and dissolution of salt-bridges with a wide range of persistence. Also, the continuous dynamic interchange of charged-atom-pairs (coming from a variety of oppositely charged side-chains) in the transient ionic bonds supports a model of dynamic flexibility concomitant with the well characterized stochastic conformational switching in these proteins. The results and conclusions should facilitate the future design of salt-bridges as a mean to further explore the disordered-globular interface in proteins.

## 1. Introduction

Recent research in Intrinsically Disordered Proteins (IDPs) has brought about a paradigm shift in the central dogma of protein folding, specifically questioning the ′one sequence one structure one function′ epitome in globular proteins [1]. IDPs lack unique three-dimensional structures under physiological conditions, but are known to carry out a multitude of important biological functions. The intrinsic disorder of a protein may be classified as i) Intrinsically Disordered Protein Regions (IDPRs) where well-defined secondary structures coexist with highly flexible disordered regions and ii) Intrinsically Disordered Proteins (IDPs) which are completely devoid of any distinct tertiary structure. The functional diversity of IDPs/IDPRs may be attributed to their potential promiscuity in binding to different structurally unrelated partners in living cells [2]. IDPs and IDPRs are usually represented as dynamic ensembles of interconvertible conformations [3] instead of well-defined structures that characterizes globular proteins. The structural flexibility of IDPs/IDPRs [4] in uncomplexed form imparts higher conformational entropy [5] due to continuous fluctuations between multiple conformers. Compared to globular proteins, IDPs and IDPRs are highly enriched in charged and polar residues and deficient in hydrophobic residues, which is denoted by a higher mean net charge to mean hydrophobicity ratio [2,6,7]. It has been observed in protamines (a class of arginine-rich IDPs with defined cellular functions) that the net charge per residue is one of the discriminating ′order′ parameters responsible for the adoption of heterogeneous conformational ensembles in IDPs/IDPRs during the globule-to-coil transition [8]. Also, in contrast to the globular proteins, the hydrophobic residues are randomly scattered along the disordered sequences [4,9] that do not mediate a hydrophobic collapse [10,11] accompanied by concomitant water depletion.

Theoretical studies based on designed globular and disordered protein sequences have revealed that there is a transition of the prevalent interactions at the globular-disorder interface across the charge-hydrophobicity boundary [12]. To this end, the globular-disorder (or globule-to-coil) transition may be explained based on a compromise between hydrophobic and electrostatic interactions. The prevalence of polar and charged residues in the disordered proteins potentially trigger electrostatic interactions, both attractive and repulsive, which constitutes the predominant interaction in IDPs/IDPRs [13]. The role of electrostatic interactions are manifest in the binding of IDPs/IDPRs to their targets and in the binding of histone chaperones, [14] which are investigated by combined NMR and Molecular Dynamics (MD) studies [15]. One of the key cellular functions seldom observed in IDPs is their phosphorylation, initiating ultra-sensitive reactions, where the electrostatic interactions play a pivotal role, contributing to ultra-sensitivity [16]. Again, one of the major electrostatic components are the salt bridges, that comprise of ionic bonds between oppositely charged amino acids in close contact. Salt-bridges, both in isolation and as composite ionic bond motifs [17], are known to impart local rigidity in protein structures [18] (analogous to the disulphide bridges [19,20]) by bringing together distant regions in folded globular proteins, acting as molecular clips at the protein-protein interface and by promoting loop-closures as revealed in computationally modeled partially disordered proteins [17]. However, the structural plasticity of IDPs/IDPRs, rich in prolines and glycines gives rise to high coil forming propensities [4]. Hence, it would be logical to explore the dynamics of salt-bridge formation in partially/completely disordered proteins for the understanding of how IDPRs/IDPs retain structural flexibility at the expense of a large number of charged residues.

The main purpose of this study is to understand the meticulous trade-off between these two seemingly opposite phenomena which may facilitate the design of partially/completely disordered proteins. With this background, we explore the structure and dynamics of these salt-bridges, both individually and also as composite motifs in IDPs/IDPRs along their simulated MD trajectories. We analyze their dynamic persistence, their involvement in mediating secondary structural ′order-disorder′ transitions and other associated structural features in an attempt to rationalize their paradoxical presence in good numbers in the dynamic conformational ensembles of IDPs/IDPRs. We also investigate in detail, the specific contribution of these salt-bridges in depicting the overall electrostatic interaction map of these proteins, given the importance of electrostatic interactions in IDPs/IDPRs. Furthermore, we also attempt to explain the solution-reactivity of IDPs/IDPRs from the analyses of their respective hydrophobic burial profiles.

## 2. Materials and Methods

### 2.1. Selection of IDPs

To study the salt-bridge dynamics in IDPs/IDPRs, a set of four disordered proteins were chosen with varying degrees of structural disorder (ranging from 43 to 100%) in their native states, containing oppositely charged amino acids (roughly 1/3^rd^) ensuring the formation of salt-bridges (**Supplementary Table S1**). The set consisted of two partially disordered proteins (IDPRs); i) the scaffolding protein GPB from Escherechia virus phix174, PDB ID 1CD3, chain ID: B, ii) the human coagulation factor Xa, PDB ID: 1F0R, chain ID: B and two completely disordered proteins (IDPs), namely α-synuclein (α-syn) and amyloid beta (Aβ_42_). Amyloid beta with its exact sequence length of 42 residues (Aβ_42_) was chosen. Both IDPRs, 1CD3 and 1F0R consist of long contiguous stretches of disordered regions characterized by missing electron densities in their respective PDB files. These disordered stretches in IDPRs are located at the N-terminus for 1F0R, while for 1CD3 it is mainly confined to the middle regions. The sequences of the IDPs were obtained from the DISPROT database [21].

### 2.2. Atomic Model Building

The X-ray crystallographic structures of both 1CD3 and 1F0R (resolution: 3.5 Å & 2.1 Å and R-factor: 0.275 & 0.216 respectively) have long contiguous stretches of missing coordinates (i.e., missing electron densities) corresponding to the disordered regions. These missing disordered residues were identified by comparing the SEQRES and ATOM records in their corresponding PDB files and cross-checked with those declared in the REMARK 465 list. The disordered regions for the partially as well as completely disordered proteins were modeled using MODELLER [22]. The modeling of the disordered regions is done in a manner such that it exactly preserves the structure of the ordered part of the protein, i.e., the RMSD (root-mean-square deviation) of the ordered part of the protein is exactly zero. Such methods were followed in earlier works [5,6].

### 2.3. Molecular Dynamics Simulation

Explicit-water Molecular Dynamics (MD) simulation trajectories for 1CD3, 1F0R and α-syn used in the current calculation were directly obtained from a previous study [6] while, for Aβ_42_, an identical simulation protocol was followed [6]. The simulations were performed with AMBER 12 program [23] at T=300 K using the ff99SB force field [24,25] with periodic boundary conditions and TIP3P water model [26]. Energy minimization of each solvated protein was performed twice via the conjugate gradient method to remove unfavorable steric interactions. The energy minimized solvated protein was equilibrated in an NVT ensemble for 100 ps at an initial temperature of 100 K, while the temperature was gradually raised to 300 K at constant volume. This was followed by NPT equilibration for 5 ns at a constant temperature of 300 K and a pressure of 1 bar. An NPT production run of 100 ns with a time step of 2 fs was performed on the equilibrated system of each protein. Berendsen′s temperature bath was used to maintain constant temperature with a coupling constant of 2 ps, while constant pressure was regulated by a barostat with a coupling constant of 1 ps. Trajectories were written at an interval of 2 ps, resulting in 50000 frames (or snapshots) and all analyses were performed on the post-equilibrium 100 ns long trajectories for all four proteins.

### 2.4. Globular Protein Database

A database of high resolution X-ray structures of globular proteins were compiled as a reference, using the advanced search protocol of PDB [27] with the following culling criteria: (i) resolution ≤ 2 Å (ii) neither working nor free R-factor worse than 0.2 (iii) files containing only uncomplexed proteins without DNA / RNA / DNA-RNA hybrid, (iv) a sequence identity of maximum 30% (v) only wild type proteins including expression tags and (vi) percentage coverage of sequence 90% (Uniprot) [28]. The application of the above culling criteria resulted in 2777 unique chain entries which are mapped to 2692 PDB structures (**Supplementary Dataset S1**). The resulting database is referred to as GDB and was used as a benchmark to assess and evaluate certain parameters related to IDPs/IDPRs for the first time, to the best of our knowledge.

### 2.5. Secondary Structural Assignments

Secondary structures (helices, strands, coils etc.) were determined from the atomic coordinates by STRIDE [29] and assigned to each amino acid residue in the polypeptide chain based on the available knowledge of both hydrogen bonding pattern and the backbone geometry of existing protein structures in the PDB.

### 2.6. Identifying salt-bridges

In accordance to a recent study [17], ionic bonds within disordered proteins were detected when a positively charged nitrogen atom from the side-chains of lysine (NZ), arginine (NH1, NH2) or positively charged histidine (HIP: ND1 NE2, both protonated) were found to be within 4.0 Å of a negatively charged side-chain oxygen atom of glutamate (OE1, OE2) or aspartate (OD1, OD2).

### 2.7. Salt-bridge Motifs

The construction of salt-bridge networks (also known as ‘salt-bridge motifs’) was extensively discussed in a previous study [17]. Briefly, in these networks, inter-connected charged residues were represented as nodes and the ionic bonds between them as edges. Each network (or motif) was then represented numerically describing its topology. This numerical string called the ′motif identifier′ is a concatenated string of numbers (separated by a delimiter ′-′) where each number-string stands for the topological description of a node in the network; the first digit in the number-string refers to the degree of that node, and the other digits refer to the degrees of its direct neighbors, sorted in a descending order. These concatenated number strings are then represented as elements of an array, further sorted in descending order, and concatenated by a delimiter ′-′. The formulation of the motif identifier is based on the assumption that the degrees of such nodes will be restricted to one digit numbers (i.e., a maximum of 9) -which holds true for a wide plethora of contact networks among proteins [33] including ionic bond networks [17]. A pictorial illustration of the numeric scheme can be found in Figure 2 of the original work [17].

### 2.8. Dynamic Persistence of salt-bridges

To calculate the coordinate-based dynamic properties of salt-bridges, simulated conformations were examined at a time interval of 50 ps across the entire 100 ns simulation trajectory for each protein. This resulted in a total of 2000 protein conformations assembled from different time frames. The dynamic persistence *(pers*) of a particular salt-bridge was calculated as the ratio of the number of protein conformations containing the salt-bridge with respect to the total number of conformations examined. Persistence was calculated for all ionic bonds that were formed at least once in the entire simulation trajectory. An optimum persistence cut-off of at least 25% (i.e., *pers ≥* 0.25 : a salt-bridge found in at least ¼^th^ of all sampled frames) was considered (see section **Persistent salt-bridges** in **Results and Discussion**) to filter out the most relevant ′persistent′ salt-bridges. To check the consistency of the results, the entire calculation was repeated at smaller (20 ps) and larger (100 ps) time intervals, as compared to the original time interval of 50 ns. The deviation in the corresponding persistence values were found to be negligible, as reflected in the calculated pairwise root mean square deviations (0.0035: 20 ps vs. 50ps; 0.0051: 100 ps vs. 50 ps) (**Supplementary Table S2**) of the persistent salt-bridges. The choice of the 50 ps time interval was motivated by the range of reported time scales for secondary structural transitions in proteins [31].

### 2.9. Contact Order

Contact Order (CO) of the interacting pairs of ions involved in a salt-bridge was determined as the number of residues that separates any two charged amino acids in the sequence space of the respective amino acids divided by the full length of the polypeptide chain as reported in an earlier study [17]. The analysis of the distribution of CO-values for the salt-bridges are guided by a previous work [17]. In contrast to the established use of CO as an index of the refolding rates in globular proteins [32], this measure was used for the disordered proteins to investigate whether the ionic bonds are able to bring distant parts of the polypeptide chain in close proximity; and, if so, how persistent are such long-range ionic bonds in IDPRs/IDPs.

### 2.10. Combined Disorder Status for Persistent salt-bridges

One of the major goals of this work is to depict the role of salt-bridges in the dynamics of conformational interconversions in IDPs/IDPRs. To this end, each charged residue involved in a salt-bridge, in a given time frame, was categorized as either ‘Disordered (D)’ or ‘Ordered (O)’ according to its assigned secondary structure (STRIDE [29]) combined with the following classification scheme. The ′ordered′ class consisted of α-helices, β-strands, 3_10_-helices and the rare occurrences of П-helices while the disordered class comprised of turns, coils and bridges. Hence, the ‘combined disorder status’ (CDS) of two charged residues involved in a salt-bridge in a given conformation may be classified as either disordered (DD) or ordered (OO) or hybrid (DO / OD). Each possible transition that a salt-bridge may undergo in terms of this CDS between two conformations at consecutive time intervals (e.g., OO → DD, OO → DO etc.) may be assigned a score based on how many of the charged residues involved in the salt bridge pair actually undergo the transition. In accordance, the following scoring scheme (**Supplementary Table S3**) was adapted. Special care was taken to assign a score resulting from the swapping of hybrid CDS′s (i.e., OD→ DO and vice-versa).

These scores (or transition weights) for a particular pair of charged residues in a salt bridge were then summed for the conformations at two consecutive time intervals along the simulation trajectory and divided by the total number of such transitions, generating the conservation score for salt-bridge secondary structures (CS_sec_). Actually, CS_sec_ is a discrete probability measure defined in the range of 0 to 1; where 0 and 1 refers respectively to cases of none and both residues in the pair undergoing changes in their respective disorder-status.

### 2.11. Shape Factor

Shape factor (ρ) is a well known descriptor of shape and compactness for polymers. This parameter (also known as internal structure factor) approaches a value of 1.5 and 0.77 for an ideal chain (also called random coil) and a compact sphere respectively [33–36]. Shape factor may be calculated from the atomic coordinates [37] as the ratio of the radius of gyration (R_g_) and the hydrodynamic radius (R_hyd_), where the two radii, R_g_, R_hyd_, are defined by the following expressions:

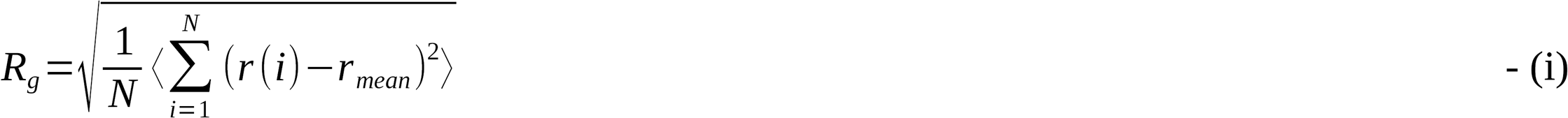

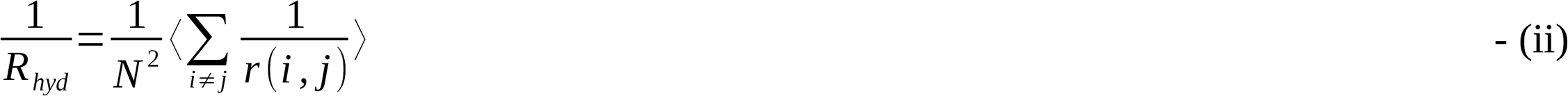

where r(i) is the distance of the i^th^ atom from the centroid of the molecule, denoted as rmean and r(i,j) is the distance between the i^th^ and the j^th^ atoms in the polymer. Thus, R_g_ may be defined as the root mean square deviation of all atoms from the centroid of a molecule which is then divided by Rhyd to nullify the size-effect, so that, ρ may be compared across different sizes of polymers. Shape factor was calculated for different conformations of IDPs/IDPRs to explore the dynamic shape transition of these proteins along the simulation trajectory due to the salt-bridge dynamics and other electrostatic interactions.

### 2.12. Burial of Solvent Exposure

The solvent accessible surface area (ASA) was calculated for each protein atom by NACCESS [38] as in earlier studies [17,39]. The ASAs were summed up for all atoms for a given residue. The extent of burial *(bur)* or solvent exposure for a specific residue X was calculated as the ratio of the ASA of X located in the protein to the ASA of the same amino acid in a Gly-X-Gly peptide fragment in its fully extended conformation [40]. Residues were distributed according to the extent of burial or solvent exposure with empirical boundaries standardized in earlier works [17,30,40]. Precisely, four burial bins were considered that quantifies the degree of solvent exposure: bur1: 0.0 ≤ *bur* ≤ 0.05 (completely buried); bur2: 0.05 < *bur* ≤ 0.15, bur3: 0.15 ≤ *bur* ≤ 0.30 (partially buried) and bur4: *bur* > 0.30 (completely exposed).

### 2.13. Accessibility Score

The accessibility score *(rGb)* is a knowledge-based measure that compares the hydrophobic burial profile of a given protein with respect to those of the highly resolved globular protein structures. The empirical ranges of this measure have been standardized on globular proteins [41] and also at protein-protein interfaces [42] and they seem to unambiguously agree with each-other. It may be defined as follows.

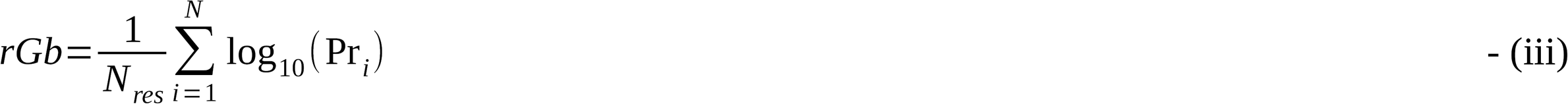

where N_res_ is the total number of residues in the polypeptide chain, Pr_i_ is the propensity of the *ith* amino acid (Ala, Glu, Asp etc.) to acquire a particular degree of solvent exposure. Other details of the score is defined elsewhere [41]. The score is a strong indicator of stability or instability of a protein / peptide in solution while instability may be further extended to explain reactivity.

### 2.14. Electrostatic Complementarity and Score

Electrostatic complementarity (E_m_) for amino acid residues in all proteins were calculated precisely following previous studies [17,43] and a modified version of a log-odd probability score (P_Em_) was computed for the whole protein chain as an overall descriptor of electrostatic interactions adapted from a previous formulation [41]. Briefly, the score can be defined as follows.

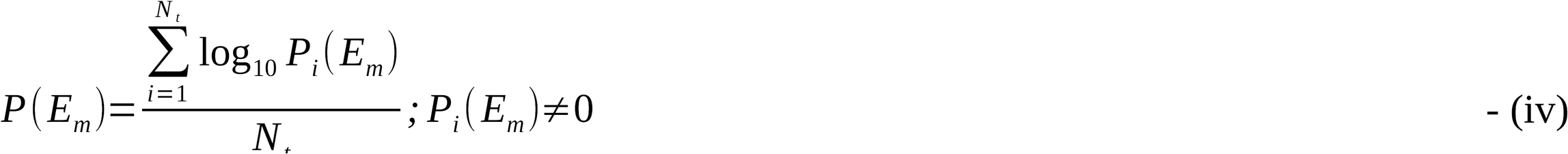

where N_t_ is the total number of amino acid residues in a given polypeptide chain and Pi(E_m_) is the probability of the i^th^ amino acid residue to acquire an Em value which falls in a particular bin of discrete probabilities (P (x < E_m_ < (x+0.05))) within the defined range of E_m_, varying from -1 to +1. Rather than considering only the completely or partially buried residues [41], all four burial bins [41,43], including the solvent-exposed residues were considered in this work. Also, as opposed to the original study [41] where P_Em_ was calculated only on E_m_^sc^ (E_m_ computed on the side-chain van der Waal′s surface of the target amino acid [43]), here, the modified P(E_m_) score was calculated for all three E_m_ measures, namely, E_m_^all^ (E_m_ calculated on the van der Waal′s surface of the entire residue of the target amino acid), E_m_^sc^ corresponding to its side-chain surface and E_m_^mc^ computed on its main-chain surface [43]. These additions have modified the representation of the score, P(E_m_) in contrast to P_Em_ [41] such that the three different E_m_ measures can be conveniently denoted as its potential argument (viz., P(E_m_^all^), P(E_m_^sc^), P(E_m_^mc^)). All force field parameters (partial charges, van der Waals radii), wherever applicable, were used in consistency with the original work [43].

### 2.15. Dynamic Cross Correlation Map (DCCM)

The Dynamic Cross Correlation Map (DCCM) [44] captures the time correlated positional fluctuations due to the concerted movement of the salt-bridge forming oppositely charged residues. This correlation map is a visual portrayal of the DCC(i,j), defined as follows.

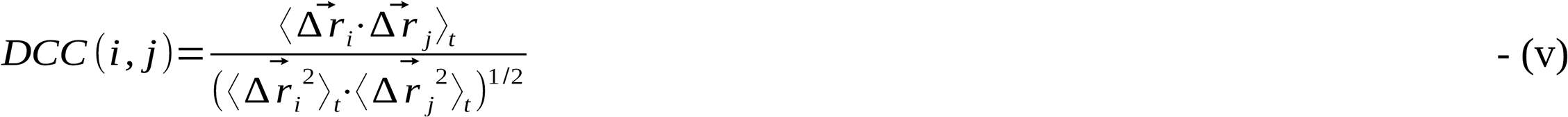

where i and j refer to the reduced single atomic representations (as defined below) of any two charged residues (taken pairwise from a pool of all charged residues) in the molecule. The displacement vector of the i^th^ residue from its mean position at the t^th^ snapshot of the simulated trajectory is denoted by Δri (i.e., its representative atom). The map was constructed respectively for both main-and side-chains of each protein. For the main-chain, the centroid of the molecular unit of Ni-C^α^_i_-C_i_ was considered as the representative of the i^th^ residue while for the side-chain, the charged atoms in the ionic groups were considered. For Lysine, the only charged side-chain atom NZ is considered, while for residues with bifurcated ionic groups involving two degenerate charged atoms of the same type (N/O), the mean position of the two charged atoms were considered (i.e., ARG: (NH1+NH2)/2; GLU: (OE1+OE2)/2; ASP: (OD1+OD2)/2; HIP: (ND1+NE2)/2). For a given protein, all structures obtained from a given simulated trajectory were first superposed on a common template to bring the coordinates on the same frame. After superposition, the time averaged mean positions of all atoms and those of the relevant atomic units were computed, followed by the calculation of the displacement vectors for each representative atom (as in equation (v)) at every sampled (t^th^) snapshot in the trajectory. 2000 equispaced snapshots were sampled at an interval of 50 ps covering the entire 100 ns trajectory. The dot product of the displacement vectors Δr_i_ and Δr_j_ at each (t^th^) snapshot were then computed, normalized by the product of their individual dot products. This ′normalized dot product′ term was then time-averaged over all snapshots in the simulated trajectory giving DCC(i,j). Hence, the values of DCC(i,j) varies between -1 to 1; wherein, -1 and 1, respectively represents perfect anti-correlation and perfect correlation between the movements of the i^th^ and the j^th^ atoms, sustained throughout the entire dynamic trajectory; while a DCC(i,j) of 0 means no correlation. Although the DCC(i,j) matrix consists of entries corresponding to all charged residues in each protein, we are only interested in the oppositely charged pairs forming the persistent salt-bridges.

### 2.16. Quantification of Hydrophobic Clustering

To analyze the contribution of hydrophobic clustering in the shape variations of IDPs, we carried out a thorough investigation of completely or partially buried hydrophobic contact networks, according to a previously established methodology [45]. Briefly, all hydrophobic residues (Ala, Val, Leu, Ile, Phe, Tyr, Trp, Met) which were either completely or partially buried within the protein interior *(bur* ≤0.30) were identified and contact networks were constructed based on the topology of their mutual adjacency. If any two heavy atoms from two different buried hydrophobic side-chains were found within a stringent cutoff of 3.8 Å^1^ from each-other, these two residues were said to be in contact. This cutoff could be treated as optimal since the resultant point atomic contact networks resembled maximally with the corresponding ‘geometrically constrained’ surface contact networks constructed from the shape complementarity and overlap between the interacting side-chain van der Waals surfaces [45]. Thus, two such buried hydrophobic residues found to be in contact served as two nodes in the above mentioned contact network and were connected by a link. To measure the extent of hydrophobic clustering, clustering coefficients (Clcf) for each individual node in these networks were then calculated according to the following expression.

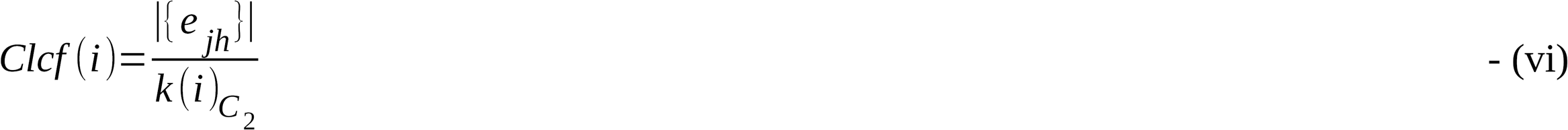

where k(i) is the degree of the i^th^ node and |{ejh}| is the total number of actually existing links among the set of nodes (taken pairwise, {j,h}) from the direct neighborhood of node i and ^k(i)^C_2_ is the (theoretical) maximum number of possible connections within the same set [46]. Values of Clcf(i) were then averaged over all nodes within a network (<Clcf>) giving the average clustering coefficient of the network. Clustering is a measure of *‘local cliquishness of a typical neighborhood’* [46] and Clcf has a theoretical range of [0,1]. That is to say that Clcf will attain a value of 1 for complete graphs^2^ or sub-graphs (cliques). Any non-zero value of Clcf is indicative of some clustering; higher the value, better the clustering. The same measure was computed in the globular protein database (GDB) to set appropriate benchmarks. The whole calculation was repeated with a slightly relaxed cutoff of 4 Å, just to reveal the scenario when an identical contact criteria to that of the salt-bridges were used. The time averages in <Clcf> were computed for all four IDPs / IDPRs and compared with corresponding values, as obtained from GDB.

### 2.17. Molecular Docking

To probe the plausible functional role of salt-bridge dynamics in IDPs, the binding of α-synuclein and tubulin [47] was chosen as a model case study, and, molecular docking was performed between the two. The whole 100 ns trajectory of α-synuclein was equally divided into 5 epochs of length 20 ns each (i.e., 0-20 ns, 21-40 and so on…) and 20 equidistant frames (at an interval of 1 ns) were sampled in each epoch to serve as receptor-templates in 20 independent docking exercises. The human tubulin coordinates solved at 2.55 Å by X-ray were extracted from the PDB ID: 4YLR and used as the ligand.

Autodock (version 4.2) [48] was used for docking, both in its GUI as well as in a customized command-line mode, as per requirement. Hydrogen atoms of tubulin were geometrically built by REDUCE [49]. Note that the receptor MD frames already had hydrogen atoms. Partial charges were assigned to all atoms (both receptor and ligands) using the ‘Add Kollmann charges’ module of Autodock, prior to docking. For the initial ‘autogrid’ settings, the grid boxes were visually set, centered on the receptor-centroid, large enough to cover all atoms of the whole receptor molecule with the dimensions of 126 Å × 126 Å × 126 Å. The ligand (tubulin) was input without opting for the ‘open as rigid’ option and thereby automatically implementing the ‘build torsion tree’ option of Autodock wherever applicable, allowing for enough flexibility for the ligand during docking. Independent *ab-initio* dockings were then performed for each ligand using a meta-heuristic GENETIC algorithm with its suggested default settings (population size 150, a maximum of 2500000 fitness evaluations, 27000 generations, gene mutation rate: 0.02, rate of cross-over: 0.8) for a run of 100 cycles. The resultant 100 docked poses were then ranked according to binding energy (lower the better) and the lowest energy conformer was selected from each cycle for further analyses.

### 2.18. Design of salt-bridge mutants in Aß_42_

To address whether the native sequence is indeed the key determinant of the salient dynamic features of the salt-bridges, a case study was performed comprising of short (30 ns) simulations performed on two appropriately chosen mutants of Aß_42_ (Aß_42_.M1, Aß_42_.M2). Four ‘XA → la’ point mutations were performed on each case (X: a selected charged residue). Alanine, was chosen as the mutated residue in all instances for three reasons: (i) its non-bulky, chemically inert, methyl functional group, (ii) to avoid the possible formation of other non-specific salt-bridges which might have occurred between mutations involving oppositely charged amino acids (say, Lys → Glu), and, (iii) in absence of a well defined hydrophobic core in the protein, issues like incorporating packing defects or holes by the choice of the small side-chain of alanine are ruled out.

In **Aß_42_.M1**, all high persistence (pers ≥ 0.25) salt-bridge forming residues were mutated to alanines

These residues were: 3-Glu, 5-Arg, 11-Glu and 13-Hip. There were two such salt-bridges to be dismantled due to this set of mutations. In the other mutant, **Aß_42_.M2**, residues forming multiple transient salt-bridges (again, four of them, to keep consistency with **Aß_42_.M1**) were mutated (residues: 1-Asp, 6-His, 16-Lys, 28-Lys), effectively dismantling 10 transient salt-bridges. Following are the sequences with the mutated positions in the mutants are highlighted in bold.

> **Aß_42_**

DAEFRHDSGYEVHHQKLVFFAEDVGSNKGAIIGLMVGGVVIA

> **Aß_42_.M1**

DAAFAHDSGYAVAHQKLVFFAEDVGSNKGAIIGLMVGGVVIA

> **Aß_42_.M2**

AAEFRADSGYEVHHQALVFFAEDVGSNAGAIIGLMVGGVVIA

Short MD Simulations of length of 35 ns were carried out using an identical protocol implemented for all native IDPs (see section. 2.3. **Molecular Dynamics Simulation**). The initial 5 ns trajectories were discarded and the post-equilibrium 30 ns trajectories retained in both mutants. All post-simulation analyses were carried out in reference to the corresponding patch of simulation trajectory (i.e., the initial post-equilibrium 30 ns) for the native Aß_42_. Frames were sampled at 25 ps interval.

## 3. Results and Discussion

IDP sequences are often abundant in polar and charged residues. These charged residues may carry like or opposite charges and may accordingly modulate the overall electrostatic balance of the protein and its binding. Salt-bridges (or ionic bonds) are the major electrostatic components involving the -NH3+ and COO-groups of the charged amino acids. However, formation of dynamically persistent salt-bridges in IDPs, especially those having long range contacts, should, in principle be a deterrent to their intrinsic flexibility, imparting local structural rigidity, analogous to the disulphide bridges in context of globular proteins. Thus there is a subtle balance between the optimum dynamic flexibility in the IDPs/IDPRs and an allowed set of salt-bridges (out of all possible pairwise combinations of residues carrying opposite charges) such that these proteins are able to accommodate the charged residues without compromising their dynamic flexibility. In this work, we investigate this trade-off in both partially and completely disordered proteins by analyzing the time evolution of the salt-bridge motifs (see **Materials and Methods**, section 2.7. **Salt-bridge Motifs**) along their simulated MD trajectories.

### 3.1. Distribution of salt-bridge motifs in the dynamic trajectories of IDPs

Composite salt-bridge motifs were identified and accumulated (see **Materials and Methods**, section 2.7. **Salt-bridge Motifs**) along the 100 ns MD simulation trajectories for all four IDPs/IDPRs. 2000 equispaced MD frames sampled at 50 ps were collected and analyzed to obtain a distribution that revealed the preponderance and insignificance of the different salt bridge motifs. Essentially we estimate the frequency of occurrence of each salt bridge motif along the simulated trajectories of each protein. The majority of salt-bridge motifs were found to be isolated (motif identifier: 11-11), amounting to 84.8%, 80.2%, 89.9% and 98.8% for 1CD3, 1F0R, α-syn and Aβ_42_ respectively (Figure 1) similar to the earlier findings on globular proteins, modeled structures of partially disordered proteins and protein-protein interfaces [17]. In accordance with increasing complexity, the second most frequent motif was found to be bifurcated salt-bridges (211-12-12) accounting for 7.7%, 15.9%, 8.9% and 1.2% in these four proteins respectively. The frequency of these motifs gradually decreased with increasing size and/or complexity of the networks. Other noticeable low-frequency motifs constituted of (i) four-membered open linear chains or trees (221-221-12-12: 5.8% in 1CD3, 1.6% in 1F0R, 1.1% in α-syn), (ii) four-membered closed rings also known as 4-cycles (222-222-222-222: 1.4% in 1CD3) and (iii) open motifs with four nodes including a central node, also known as 3-stars (3111-13-13-13: 1.8% in 1F0R). The rest of the motifs mostly containing greater than 4 nodes were found in abysmally low frequencies. The partially disordered proteins displayed an overall higher frequency of salt-bridges as revealed by the difference in the total number of motifs (1CD3: 13698, 1F0R: 14055) as compared to the completely disordered ones (α-syn: 8188, Aβ_42_: 3190). This difference is due to (i) the significantly smaller length of the fourth protein (~1/3^rd^ of the other three) and (ii) the lower fractional content of the charged residues in the IDPs (~28%) as compared to the IDPRs (~34%) (**Supplementary Table S1**).

**Figure 1.**
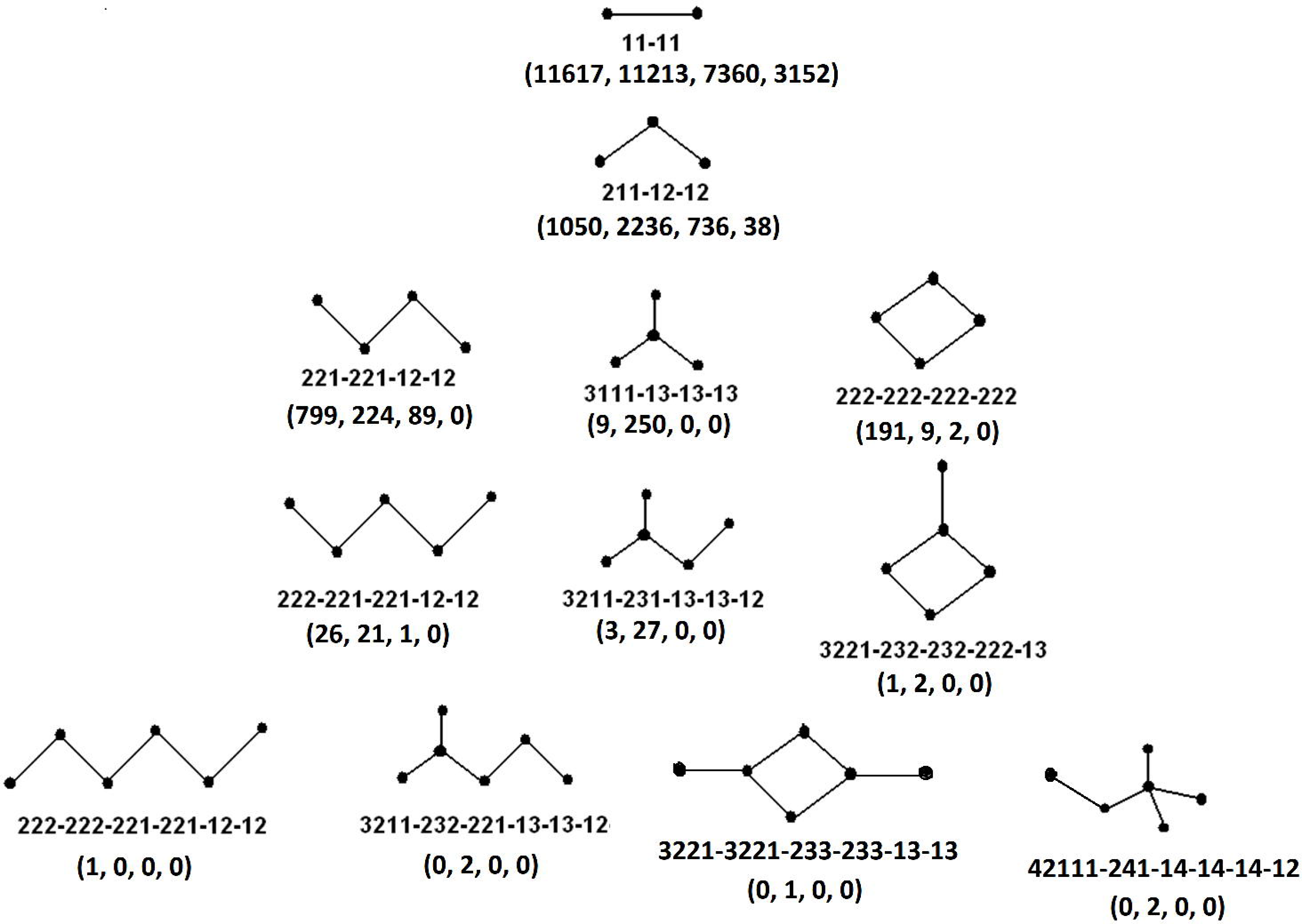
Salt-bridge motifs found in simulated conformational ensembles in IDPs. Each motif (network) is represented as a ball (node: charged residue) and stick (edge: ionic bond) model and tagged by the numeric motif identifier (see **Materials and Methods**, section 2.7. **Salt-bridge Motifs**). The numbers in parentheses stand for the cumulative counts of the corresponding salt-bridge motif in the 100 ns simulation trajectories of 1CD3, 1F0R, α-syn and Aβ_42_ respectively.

Since dynamically persistent salt-bridges counteract the flexible nature of the IDPs, imparting local rigidity, it may be interesting to find salt-bridges in IDPs/IDPRs, which may be conserved or even grow to form composite salt-bridge networks along their respective simulation trajectories. It is obvious that the salt-bridge motifs undergo transitions via formation and breakage of ionic bonds, which is evident from the analysis of the time evolution profiles (in terms of counts) of each particular motif along the MD trajectories. The length and the fraction of charged residues (**Supplementary Table S1**) of each protein influences these counts.

The number of isolated salt-bridges in both IDPRs were found to vary between 1 to 12 (mean: 5.8 ±2.1 for 1CD3, 5.7 ±1.8 for 1F0R) while for the IDPs, these numbers vary between 0 to 9 for α-syn (mean: 3.69 ±1.39) and 0 to 5 for Aβ_42_ (mean: 1.58 ±0.92) throughout the 100 ns simulation trajectory (**Supplementary Figure S1**). For the second most prevalent motif, namely, the bifurcated salt-bridges, the time-averaged counts were 0.53 (±0.63), 1.24 (±0.76), 1.22 (±0.68) and 0.02 (±0.14) for 1CD3, 1F0R, α-syn and Aβ_42_ respectively (**Supplementary Figure S2**). One interesting feature was the periodic appearance of at least one (Figure 2) and at most two (**Supplementary Figure S3**) 4-cycles or closed quadruplets without the diagonal edges (222-222-222-222), found within the 75-100 ns time frames found in all three proteins except Aβ_42_, which completely lacks the motif. Interestingly, almost all residues forming these closed quadruplets originated from the loops and coils, thereby bringing together the distant disordered regions of α-syn and the IDPRs (1CD3, 1F0R). In fact, this closed ring core topology was found in the distribution of salt-bridge motifs (Figure 1). It may be noted that the charge constraints in salt-bridges only allow for an even number of nodes in a closed ring topology (or cycle) to be found within proteins [17].

**Figure 2.**
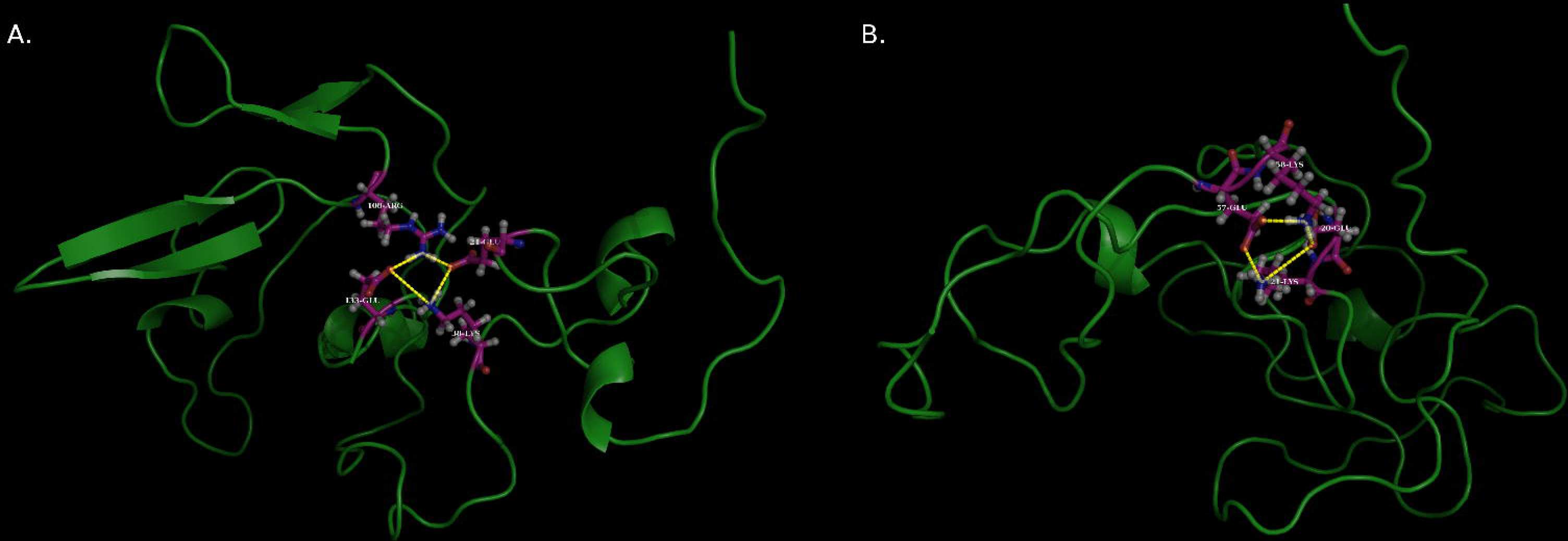
The closed quadruplet salt-bridge motif in IDPs. The 4-cycle motifs encountered in (A) 1F0R (B) α-syn within 75-100 ns MD frames. The ionic bonds are displayed as yellow dashed lines between residues represented as ball and stick models on top of the corresponding main-chains portrayed as cartoon. The residues forming the closed quadruplets are 38-Lys, 133-Glu, 21-Glu, 108-Arg (1F0R) and 21-Lys, 20-Glu, 57-Glu, 58-Lys (α-syn).

### 3.2. Persistent salt-bridges

Another pertinent problem involves the dynamic persistence of the salt-bridges — whether they are sustainable enough in the long run. To this end, the topological details of each salt-bridge motif were totally ignored and the complete distribution of the salt-bridge motifs were reconsidered as a non-redundant set of discrete ionic bonds. The dynamic persistence *(pers)* of a salt-bridge is defined as the fraction of the time-frames in which the salt-bridge / ionic bond appears with respect to the total number of time-frames sampled in the trajectory. Initially, all salt-bridges were counted that occurred at least once in the sampled time frames. These numbers were found to be 69, 102, 75 and 15 for 1CD3, 1F0R, α-syn and Aβ_42_ respectively. To assess whether the ionic bonds in these salt bridges were formed by chance, as a probabilistic consequence of the simulation protocol, their persistence values were examined. Hence, only those salt-bridges, which were formed in at least 1/4^th^ *(pers* ≥ 0.25) of the sampled time frames were screened for further analyses, accounting for slightly more than 10% of the ionic bonds. The choice of this cut-off was based on the overall frequency distribution of the persistence values (**Supplementary Figure 4A**) with a large fraction of insignificant contacts that have likely occurred by chance, detected only in a few frames (~67% of the contacts having a persistence of 5% or less).

This persistence cutoff drastically reduced the number of salt-bridges to 10, 13, 5 and 2 for 1CD3, 1F0R, α-syn and Aβ_42_ respectively by discarding the insignificant transient salt-bridges. Given the overwhelming majority of these short-lived contacts, the fluctuating dynamics of these ionic bonds may influence the local flexibility of these proteins. The calculated persistence values spanned a range from 25% to 96% with instances of dynamically stable (say, the 80%-persistent) salt-bridges in all four proteins with values hitting almost every possible 10% bin-interval in the range (**Supplementary Figure 4B**). The overall distribution also portray that the salt-bridges span different time scales ranging from instantaneous, brief, moderate and long-lived. Thus, the formation and dissolution of salt-bridges appear continuously throughout the entire simulation trajectory. This dynamics of salt-bridges supports a physical model for dynamic flexibility, which also appeared to be true visually (**Supplementary Video S1**).

The other noticeable feature, especially relevant for the ephemeral salt-bridges was the transient formation and breakage of ionic bonds associated with a particular charged atom (say, NZ of 63-LYS in 1CD3) and various oppositely charged atoms originating from different residues throughout the main-chain. This dynamic interchange of charged-atom-pairs in fluctuating ionic bonds structurally supports different alternative conformations at different temporal patches along the MD trajectory, consistent with the notion of the conformational flexibility [50] in IDPs/IDPRs. An illustrative example is shown in Figure 3 where the ε-NH3+ group of 28-Lys in Aβ_42_ has been found in contact alternatively with four different negatively charged amino acids (i.e., with their side-chain-COO-groups) at different temporal patches along the trajectory: 23-Asp (14-24 ns), 1-Asp (61-62 ns, 70-75 ns), 22-Glu (63-69 ns, 78-89 ns) and 3-Glu (97-99 ns). IDPs/IDPRs are well known for their binding promiscuity, [2,51] while here we find another level of promiscuity in them, namely, in ionic bond formation. This also supports the concept of dynamic flexibility that makes them structurally adjustable and adaptable to different conformations upon binding to different molecular partners. In a recent study, near UV-visible absorption spectroscopy have also experimentally revealed the event of ‘charge transfer’ in proteins [52].

**Figure 3.**
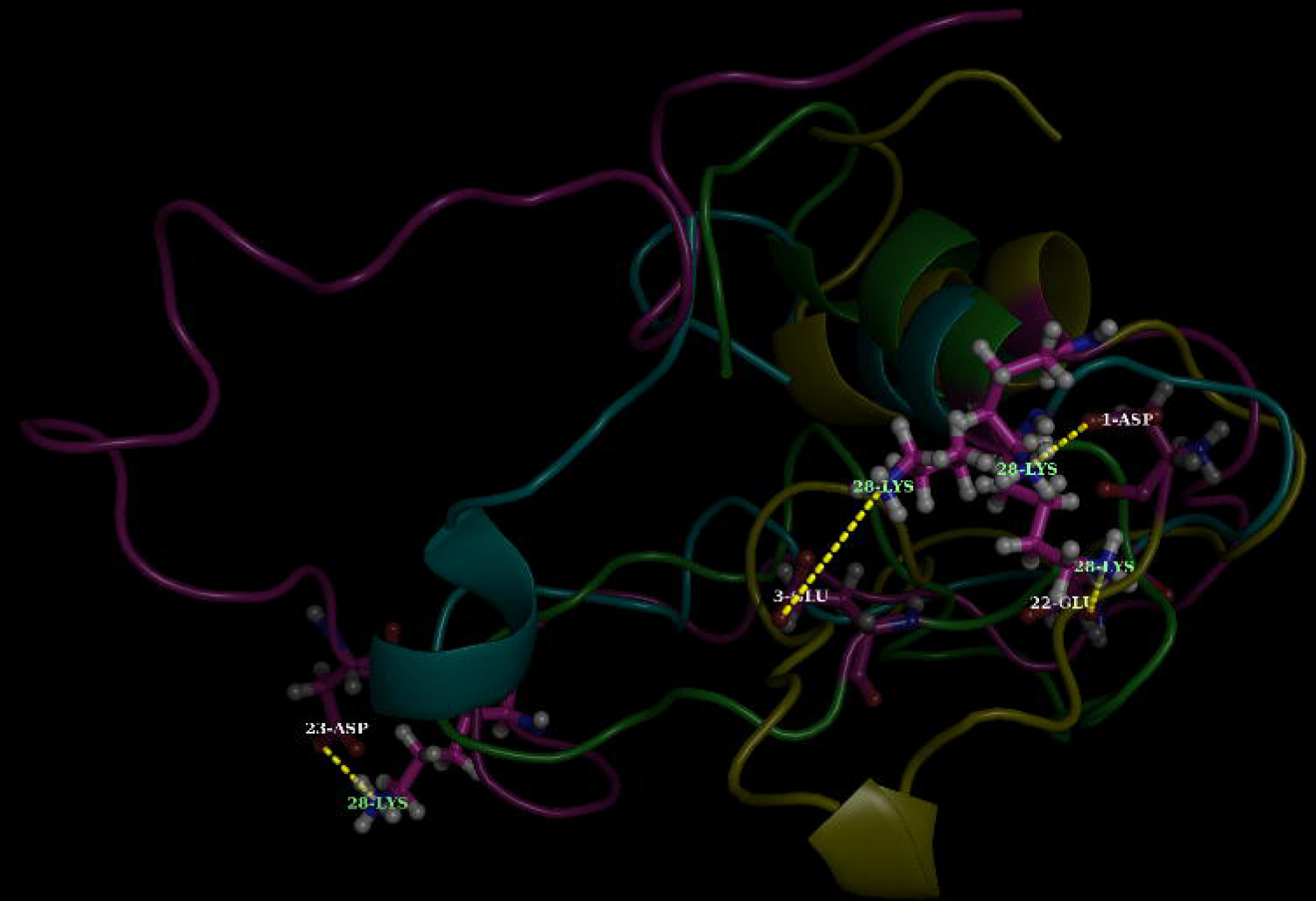
Multiple ionic bonds formed by the same charged residue at different time intervals in IDPs. Different transient ionic bonds (displayed as the yellow dashed lines) involving the amino acid 28-Lys and different oppositely charged residues (1-Asp, 3-Glu, 22-Glu, 23-Asp) are displayed at various time snapshots (see **Main Text**) in the MD trajectory of Aβ_42_. The concerned charged residues are depicted as ball and sticks in the background of the corresponding main-chains, drawn in cartoon with different colours. The different main-chain conformations were superposed on a common template prior to their display.

In parallel, it was also important to know, whether these ionic bonds were formed between residues that were close or distant in the amino acid sequence space. To that end, contact orders (see **Materials and Methods**, section 2.9. **Contact Order**) were calculated for all salt-bridges with at least 25% persistence. Most contacts (~43%) were short-ranged (Co<0.1), however, there were quite a few (~40%) middle-range contacts (0.1<Co<0.3) as well as some (~17%) long-range persistent ionic bonds (Co>0.3) too. In particular, the IDPRs (1CD3, 1F0R) contained quite a few middle and long range contacts (Table 1) and, interestingly, majority of such contacts involved an arginine (100% for 1CD3, 67% for 1F0R). Visual inspection in Pymol revealed that most contacts were susceptible to form local (short-range contacts) and non-local turns or loop-closures (long-range contacts) while some of them linked between the coils and remnants of regular secondary structural regions in partially disordered proteins, e.g., helical turns in 1CD3, beta strands in 1F0R (**Supplementary Figure S5**). For IDPs, persistent salt-bridges were exclusively found between the coils imparting some dynamic structural constraints which appeared to act against the elongated coil structure (as obtained in the initial MODELLER model) along the simulation trajectory. In α-syn, a bifurcated salt-bridge (23-LYS~28-GLU-34-LYS) was also found to be dynamically persistent even though it comprised of local contacts. In Aβ_42_, only two persistent salt-bridges were found, and both of them were on the same side of a short-helix (located towards the C-terminus of the protein and consisted of 9 residues,), which seem to gradually evolve during the course of the simulation (first appearing at around 28 ns) and found to be frequent and stable thereafter (found in 75% of the frames from 28 to 100 ns). These salt-bridges were thus structurally non-interrupting for the short helix. It is also noteworthy that except for an outwardly facing Lysine (28-Lys), the helix is otherwise devoid of any polar or charged residues; rather it consists of hydrophobic residues (Ala, Met, Leu, Ile) and Glycines only. The other interesting fact was that all charged residues (including those that does not get involved in dynamically persistent ionic bonds) were exclusively found towards the N-terminus of this short-helix, leaving it free and unperturbed (Figure 4). The secondary structures in the snapshots were depicted by STRIDE [29] and confirmed visually by Pymol and VMD [53].

**Figure 4.**
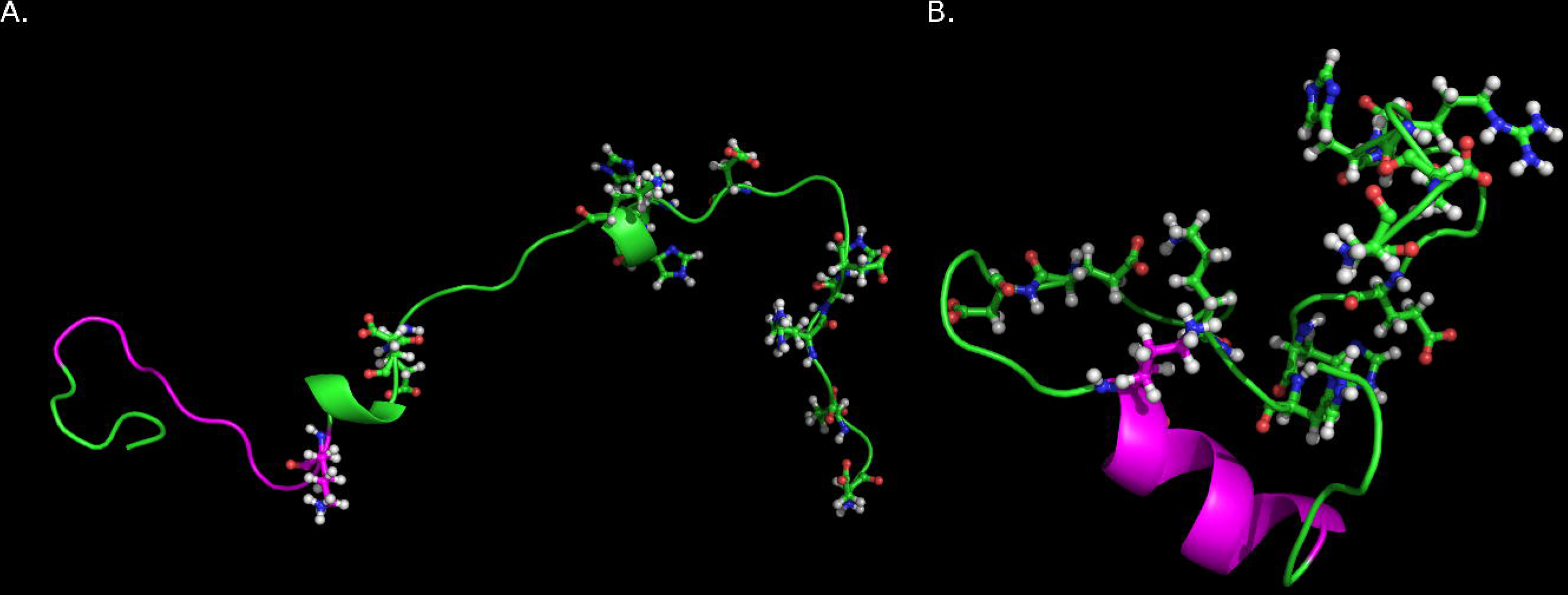
Illustration of a ‘disorder-to-order’ secondary structural transition in IDPs, as revealed in beta-amyl. The figure highlights (B) a short-helix (coloured magenta) formed during the MD-simulation run in Aβ_42_ (see **Main-Text**) and (A) the corresponding segment (same color) found as a disordered coil in the initial template structure for MD simulation. Charged residues have been portrayed as ball and sticks in both.

**Table 1.** Persistence values *(pers*) and Contact Orders (CO) of the persistent salt-bridges in the four IDPs/IDPRs.

| Salt-bridge |  | *pers* | CO |
| --- | --- | --- | --- |
| **1CD3** |  |  |  |
| 98-ASP | 100-ARG | 0.963 | 0.017 |
| 2-GLU | 108-ARG | 0.819 | 0.883 |
| 64-ARG | 68-GLU | 0.797 | 0.033 |
| 26-ARG | 28-GLU | 0.580 | 0.017 |
| 15-GLU | 50-ARG | 0.536 | 0.292 |
| 22-GLU | 76-ARG | 0.449 | 0.450 |
| 22-GLU | 64-ARG | 0.368 | 0.350 |
| 61-ARG | 65-ASP | 0.322 | 0.033 |
| 27-ASP | 77-ARG | 0.250 | 0.417 |
| 77-ARG | 105-GLU | 0.250 | 0.233 |
| **1F0R** |  |  |  |
| 90-ASP | 117-LYS | 0.818 | 0.202 |
| 28-ASP | 31-LYS | 0.803 | 0.022 |
| 10-ARG | 27-GLU | 0.726 | 0.127 |
| 10-ARG | 30-ASP | 0.643 | 0.149 |
| 14-GLU | 23-ARG | 0.448 | 0.067 |
| 21-GLU | 38-LYS | 0.430 | 0.127 |
| 82-LYS | 97-GLU | 0.401 | 0.112 |
| 108-ARG | 133-GLU | 0.378 | 0.187 |
| 15-GLU | 23-ARG | 0.335 | 0.060 |
| 41-ASP | 55-LYS | 0.321 | 0.105 |
| 96-HIP | 98-GLU | 0.311 | 0.015 |
| 58-ASP | 108-ARG | 0.285 | 0.373 |
| 62-GLU | 81-ARG | 0.267 | 0.142 |
| **α-syn** |  |  |  |
| 96-LYS | 104-GLU | 0.937 | 0.057 |
| 21-LYS | 57-GLU | 0.509 | 0.257 |
| 23-LYS | 28-GLU | 0.398 | 0.036 |
| 28-GLU | 34-LYS | 0.379 | 0.043 |
| 20-GLU | 58-LYS | 0.257 | 0.271 |
| **Aβ<sub>42</sub>** |  |  |  |
| 3-GLU | 5-ARG | 0.878 | 0.048 |
| 11-GLU | 13-HIP | 0.266 | 0.048 |

### 3.3. Salt-bridge mediated Secondary Structural Transitions

To decipher the contribution of salt-bridge dynamics in the time evolution of the secondary structural transitions, a measure involving the combined disorder status (CDS) of persistent salt-bridges was used (see **Materials and Methods**, section 2.10. **Combined Disorder Status for Persistent salt-bridges**). This parameter, namely, the conservation score for salt-bridge secondary structures (CSsec) takes into account both the number as well as the type of transition. CSsec will give zero for no transitions (i.e., 100% conservation) and will increase with increasing number of transitions. Also, the individual weights for each possible transition (e.g., OO → DD, OO → DO etc.) were standardized in such a manner that the maximum value of CS_sec_ is one, corresponding to a persistent salt-bridge of the non-hybrid type (i.e., DD/OO) undergoing transition in both of its residue-disorder-status between each pair of consecutive time intervals along the simulation trajectory (e.g., OO → DD → OO → DD… and so on) (see **Materials and Methods**, section 2.10. **Combined Disorder Status for Persistent salt-bridges**). The relative occupancy of a predominant secondary structural motif (PSSM) was also calculated (e.g., Coil~Turn, α-Helix~α-Helix etc.) for each persistent salt-bridge as a fraction of the most frequent secondary structural motif relative to all motifs found in all time frames. For a poor conservation score (corresponding to a salt-bridge more or less conserved in its secondary structural status along time), this PSSM value must be mapped to its major secondary structural motif.

For the partially disordered proteins, the conservation scores (CSsec) for the persistent salt-bridges spanned across the entire range from 0 to 1 (**Supplementary Table S4**). Though, most of the persistent salt-bridges were highly conserved in terms of their ‘combined-disorder-status’, there were some highly variable (98-ASP~100-ARG: 0.80, 22-GLU~64-ARG: 0.77 in 1CD3; 82-LYS~97-GLU: 0.99 in 1F0R) salt-bridges as well. All highly variable salt-bridges were found to contain regular secondary structural elements. Also, these high-CSsec salt-bridges correspond to low-to middle-range contacts that spans across a broad range of persistence values (0.37 to 0.81). No correlation was found between the *pers* and CS_sec_ and hence they appear to be independent dynamic properties of these proteins. The predominant secondary structural motif for the salt-bridges in 1CD3 involved both structured as well as unstructured coiled regions, whereas, those for 1F0R were mainly unstructured.

For the completely disordered proteins, all persistent salt-bridges were highly conserved in their ‘combined-disorder-status’ except one particular salt-bridge: 28-GLU~34-LYS (CS_sec_: 0.76) where the predominant secondary structural motif involves a rare 310-Helix (PSSM: 0.754). This is expected since these proteins rarely involve ordered regions along the simulation trajectory and due to the fact that the above discussed (see section 3.2. **Persistent salt-bridges**) short-helix formed in Aβ_42_ (Figure 4) is mostly devoid of any charged residues (but for the terminal Lysine: 28-Lys) thereby eliminating its scope to participate in a salt-bridge.

### 3.4. Dynamic bending in IDP chains

A protein, being a hetero-polymer, can not grow linearly for an indefinite period in space; it has to take a turn at some point [31,54]. This was the key idea behind the proposition of the cotranslational folding model in proteins [55,56], later proven experimentally [56] and supported by plausible genetic mechanisms (viz., pause codons) [57]. The model states that a protein chain begins to fold as soon as its N-terminus is exposed to the folding site from the translocation complex [55]-which is generally valid for all proteins undergoing ribosomal translation. In the current work, however, we were restricted to use atomic models, partially or completely built by MODELER, as the starting templates for the MD simulations (because of the lack of their experimental structures). These modeled structures resemble elongated random coils marked by the lack of any tertiary structures as revealed visually by Pymol. However, when the time-evolved simulated structures were displayed visually (in VMD), substantial dynamic bending with respect to the elongated initial template were evident as the electrostatic interactions overwhelmingly contributed to the persistent salt-bridges, bringing together distant regions of the protein chains (**Supplementary Video S1**). However, the curved structures changed continuously within a non-rigid (yet non-random) statistical ensemble of conformations, characterized as stochastic conformational switching in IDPs [50]. One way to quantify this bending or reshaping is to calculate the shape factors [37] which is the ratio of the radius of gyration and the hydrodynamic radius of the molecule (see **Materials and Methods**, section 2.11. **Shape Factor**). Shape factor is an indicator of the shape and compactness of polymers accounting for their dynamic shapes induced by the change of chemical environments [58]. As a benchmark, shape factors of globular proteins (ρ_glob_) were calculated from the database GDB (see **Materials and Methods**, section 2.4. **Globular Protein Database**). The ρ_glob_ values were found to be extremely stable, (characterized by a standard deviation of ~1/18^th^ of their mean value), restricted within a narrow bin of 1.0 ±0.054 (Figure 5). For calculating the shape factor in IDPs/IDPRs, the modeled structure was considered as the 0^th^ snapshot to provide a reference value followed by 2000 post-equilibrium snapshots (sampled at 50 ps interval). The value of the shape factor for the initial template structures of the disordered proteins varied between 1.3 to 1.6, close to the theoretical value of ρ for ideal chains, 1.5 [33]. The lowest ρ value (1.23) was recorded for 1CD3-a finding which is consistent with the fact that it has the highest fraction (~66.7%) of ordered secondary structure (**Supplementary Table S1**) among the four disordered proteins. These initial ρ values were found to fluctuate in the same range for a brief duration before attaining an equilibrium value of ~1.0 ±0.054.

**Figure 5.**
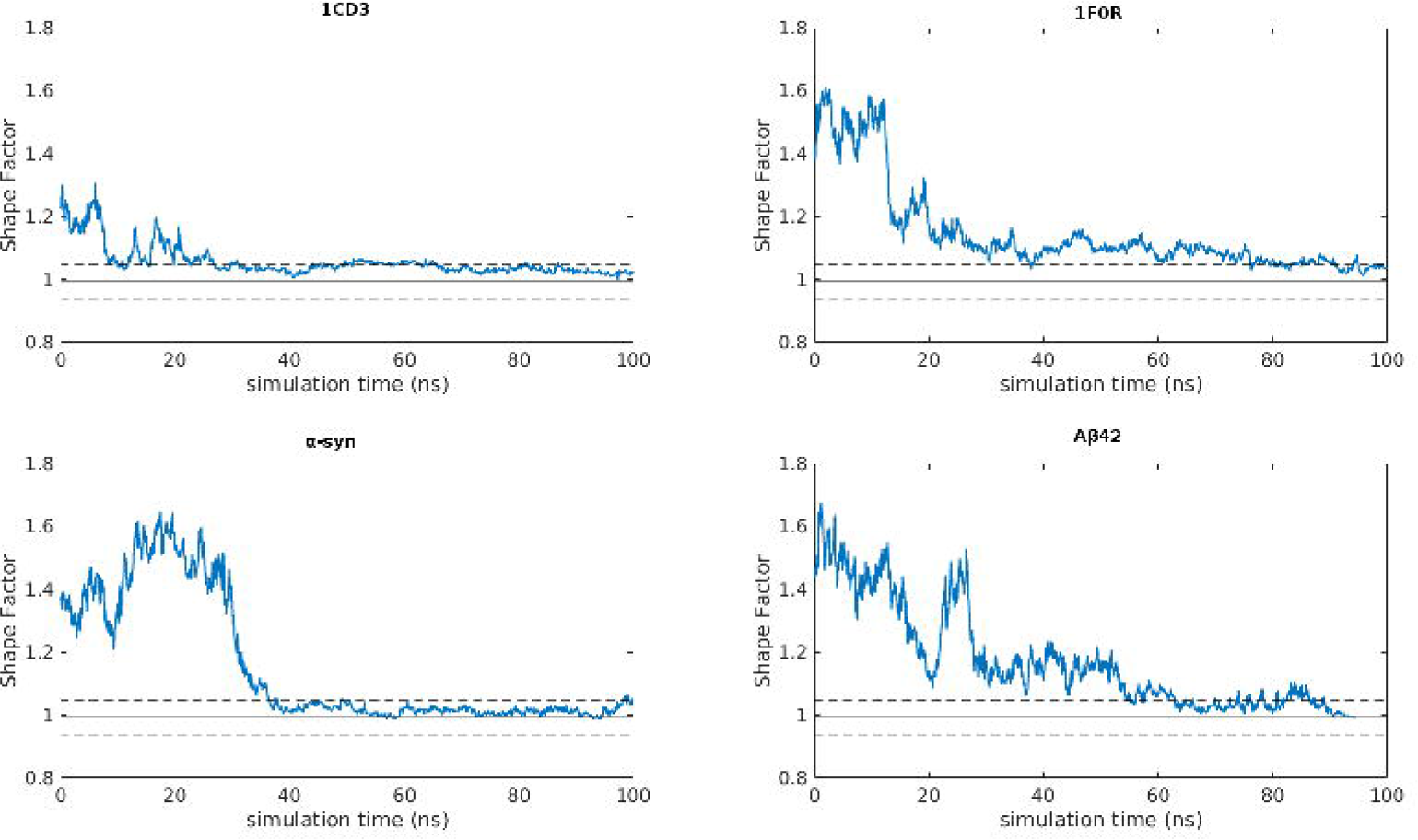
Time-evolved shape factor profiles of IDPs with respect to the globular protein reference range. Shape factor values for each individual IDP plotted against simulation time. As a reference, the mean shape factor value (p) of the globular proteins (p_glob_) is displayed as the continuous horizontal line drawn at x~1.0 flanked by two dashed lines, above and below, mapping to (μ+σ) and (μ-σ) respectively, where o stands for the standard deviation in p_glob_.

#### 3.4.1 Impact of Hydrophobic clustering, if any

Further, to study whether the dynamic bending was associated with any level of hydrophobic clustering, we thoroughly analyzed all contact networks of completely and / or partially buried hydrophobic residues (see **Materials and Methods**, section 2.12. **Burial of Solvent Exposure**), precisely according to a previously established methodology [45]. Clustering coefficients (see **Materials and Methods**, section 2.16. **Quantification of Hydrophobic Clustering**) averaged over all networks (<Clcf>) in a frame were calculated and used for further analyses. Negligible or insignificant hydrophobic clustering was observed in the completely disordered proteins (as reflected in their average clustering values, <Clcf>: 0.0293 ±0.0092 (α-syn); 0.0045 ±0.0025 (Aß_42_)) (**Supplementary Figure S6**) throughout their respective MD simulation trajectories (**Supplementary Figure S7**) compared to the values characteristic of globular proteins (obtained from the database, GDB: 0.1530 ±0.0583). Among the two partially disordered proteins (IDPRs), 1CD3 exhibited some amount of clustering (<Clcf>: 0.0396 ±0.0116: 1/4^th^ of the <Clcf> of the globular proteins) which is due to the fact that 1CD3 has the highest secondary structural content (56.7%) among all four disordered proteins. The results were virtually identical for both cutoffs (3.8 Å, 4.0 Å) (**Supplementary Figure S6**). These numbers virtually rule out the scope of hydrophobic clustering playing any appreciable role in the time-evolved dynamic bending, revealed in the shape factor profiles of the IDPs. Rather, the major component in this bending indeed appears to be the long-range electrostatic interactions, inclusive of not only salt-bridges but also hydrogen bonding, dipole-dipole and charge-dipole interactions [59]. The determining force for electrostatic interactions is essentially non-local which originates from the overall distribution of surface electrostatic potentials [43] throughout the disordered chains and can potentially bring together far-apart oppositely charged regions given certain structural contexts (e.g., in case of the high persistence long-range salt-bridges in 1CD3: 2-Glu ~ 108-Arg, 27-Asp ~ 77 Arg: Table 1). Past studies [60] have also characterized IDPs to populate compact states (or dynamically bent ensembles) due to the formation of hydrogen bonds and salt bridges, rather than to remain in extended forms all-throughout.

#### 3.4.2 Dependence of dynamic bending on chain-length and amino acid composition

To investigate the effects of distinct amino acid compositions and the chain-length in dynamic bending, shape factor profiles were regenerated for the three individual domains of α-synuclein: the N-terminal lipid-binding α-helix (1-87), the amyloid-binding central domain (NAC: 61-95), and C-terminus acidic tail (96-140) [47,61]. Overall, the general descending trend was retained in all the domains. However, both the NAC and the C-terminal domain converged (after ~ 30 ns) to a distinctly higher ρ value (ρ ~ 1.1 to 1.2) as compared to either the N-terminal domain or the full-chain (ρ ~ 1.0 to 1.05). The percentage drop with respect to the maximum ρ values ((max(ρ)-min(ρ))/max(ρ)) were 31%, 35%, 18% and 40% for the N-terminus, the NAC, the C-terminal domains and the full chain respectively (**Supplementary Figure S8**). Thus a combined effect of both domain-length and the amino acid composition is crucial for the extent of dynamic bending; larger the length and higher the number of oppositely charged residues, higher is the scope of bending. Noteworthy is the differences in length between the longer N-terminal (87 residues) and the shorter NAC (35 residues) and C-terminal domains (45 residues). Also noteworthy is that the N-terminal domain contains amphipathic repeats [47], (albeit it is more rich in Lysine; net-charge: +3) allowing for a greater degree of electrostatic attraction. In contrast, the C-terminal domain is almost purely acidic (net charge:-12) allowing little scope for the same. Note that the C-terminal domain has distinctly lower fluctuations (σ(ρ)=0.0382; σ: standard deviations) in its time-evolved ρ-values compared to the others (0.1047, 0.1632, 0.2027 for N-terminus, NAC, full-chain respectively).

### 3.5. Degree of Solvent Exposure: Approaching Porous Globules

There is very little scope for interior packing to play a pivotal role in the dynamics of IDPs/IDPRs, as they adopt expanded conformations and remain sufficiently exposed to the solvent, while undergoing continuous dynamic bending. So to speak, the packing densities of disordered proteins are expected to be remarkably different from those of the globular ones, which are similar to that of the crystalline solids [62]. There are different measures of atomic packing with different degree of details [30,37,40,63,64], however, for each and all such measures, a primary condition to retain optimum packing is to have a substantial fraction of amino acid residues deeply or partially buried within the protein interior. Thus the fraction of buried and solvent exposed residues in these IDPs were examined along the simulation trajectory. The residues were then distributed in four previously standardized burial bins [40], and the fraction of the residues present in each bin was calculated. The calculation was performed on 500 snapshots sampled at 100 ps intervals for all four IDPs. The time-averaged fraction of residues present in the four burial bins (bin1: completely buried, bin2, 3: partially buried / partially exposed, bin4: completely exposed; see Materials and Methods, section 2.12. Burial of Solvent Exposure) were 0.03 (±0.02), 0.07 (±0.03), 0.15 (±0.05) and 0.75 (±0.09) respectively for all four proteins. Clearly, the bulk share (~75-80%) of residues remain exposed to the solvent throughout the entire simulation trajectory-which was also confirmed by the actual time evolution plots (**Supplementary Figure S9**).

When the burial profiles of the IDPs/IDPRs are viewed together with their corresponding shape factor profiles-it is revealed that despite dynamic bending (as elaborated in the earlier section), the molecules still remain substantially exposed to the solvent. These features closely resemble that of the disordered collapsed globules [65,66] often encountered in redesigned proteins with over-or under-packed cores [66]. Due to packing defects, these redesigned globules become enough porous and eventually result in the complete opening up of the structures, thereby allowing water molecules to penetrate into the protein core. The water molecules entrapped this way within the collapsed globules have further been characterized as potentially important for controling crucial IDP-events like binding induced folding and amyloid aggregation, by virtue of their unique kinetic features [65]. Also, since, optimum tertiary packing can not occur in these molecules with such high degree of solvent exposure (due to the lack of enough nearest neighboring surface points to pack against), these results also effectively rule out ‘packing’ to be a major component in the overall structural feature in the dynamic ensemble of IDPs/IDPRs.

To further characterize the hydrophobic burial profile of the amino acid residues in IDPs/IDPRs (with reference to the globular proteins), the accessibility score (*rGb:* see Materials and Methods, section 2.13. Accessibility Score) was estimated for the time-evolved ensemble of structures. This parameter *(rGb)* estimates the distribution of different amino acid residues (i.e., hydrophobicities) with respect to their burial in a given structure and calibrates that structure with reference to the same estimate derived from natively folded globular proteins. However, the distribution of *rGb* is quite broad, characterized by a μ=1.5σ^3^ width both in globular proteins (μ=0.055 ±0.022) [41] as well as in protein-protein complexes (μ=0.059 ±0.022) [42] accounting for a variety of possible distribution patterns of hydrophobic burial as a function of shape and size of proteins. The time-averaged statistics obtained for *rGb* in IDPs/IDPRs registered significantly lower values as compared to that of the globular proteins (**Supplementary Figure S10**). The *rGb* value is slightly above zero for IDPRs (1CD3, 1F0R), while it is negative for the IDPs (α-syn, Aβ_42_). All values are out of range with respect to the globular protein reference and the value for Aβ_42_ is exceptionally low. This means that in IDPs/IDPRs, the trend of hydrophobic burial is actually opposite to that in globular proteins, where, most hydrophobic residues remain exposed to the solvent. While, the charged residues may, in fact, sometimes get partially buried, surrounded by enough protein atoms due to the formation of ionic bonds-more plausible for residues involved in the composite salt-bridge motifs due to their close proximity. Indeed, ~21.7% of the charged residues forming persistent salt-bridges *(pers* ≥ 0.25) were found to be at least partially buried (time-averaged burial ≤ 0.3). Conversely, 77.6% of the hydrophobic residues (viz., Ala, Val, Leu, Ile, Met, Phe, Tyr, Trp) were found to be completely exposed to the solvent (time-averaged burial > 0.30). This reversed trend in the hydrophobic burial in IDPs/IDPRs (with respect to folded globules) should substantially contribute to their high solution-reactivity, binding promiscuity [2] and the high entropic cost required to stabilize them [67]. All these features are in accordance with the ‘coupled binding and folding’ model [68-70] (also known as ′binding induced folding′ [65]) of the IDPs/IDPRs.

### 3.6. Impact of salt-bridges on the overall electrostatic balance in IDPs

Though atomic packing (or steric overlap) has been studied in IDPs, it is more relevant in the context of IDPRs [37] and has been assessed in terms of point atomic contacts, rather than surface fits as in globular proteins [30,40]. Point atomic contacts are naturally limited in portraying the true picture of atomic packing due to their inherent inability to depict the geometric match between the interacting side-chains [30]. As the unstructured regions are overwhelmingly prevalent, packing does not seem to be the major interaction required to maintain the dynamic flexibility of IDPs/IDPRs. IDPs/IDPRs does not allow hydrophobic collapse like that of folded globules. Rather, given the predominance of charged and polar residues, the global electrostatic balance is expected to be a major determinant in the formation of salt-bridges. Therefore, it is necessary to investigate the role of salt-bridges in the global electrostatic balance of IDPs. In this regard, the Complementarity Plot (CP) [41,43], an established structure validation tool originally proposed for globular proteins was used. CP analyzes the combined effect of packing and electrostatics together for amino acid side-chains embedded in proteins. This tool is built with the provision of analyzing the individual components, i.e., packing and electrostatics independently. The primary goal of this study was to analyze the role of salt-bridges in the context of the global electrostatic balance of IDPs/IDPRs. Thus the observation window was based on the calculated electrostatic complementarity (E_m_) of the amino acid residues of the disordered polypeptide chains along their MD trajectories. Hundred equidistant snapshots were collected at an interval of 100 ps to represent the dynamic ensemble of each protein. The software suite for CP (http://www.saha.ac.in/biop/www/sarama.html) returns three different measures of electrostatic complementarities, E_m_^all^, E_m_^sc^ and E_m_^mc^ (see Materials and Methods, section 2.14. Electrostatic Complementarity and Score) computed on ‘all’, ‘side-chain’ and ‘main-chain’ van der Waal’s dot surface points of the ‘target’ amino acid residue respectively. The original plots in CP uses E_m_^sc^ in accordance to the corresponding shape complementarity measure S_m_^sc^. For the current purpose, we were particularly interested in E_m_^all^ since the overall impact of the salt-bridges on the full residue surfaces was of prime interest. The log-odd probability score (P(E_m_)) (see Materials and Methods, section 2.14. Electrostatic Complementarity and Score) summed over amino acid residues distributed in different burial bins of the entire poly-peptide chains gives an estimate of its overall electrostatic balance. Only buried and partially buried residues that are present in the first three burial bins were considered in the original study [41], however, for this work, the fourth burial bin corresponding to the solvent exposed residues *(bur* > 0.30) were also considered (see Materials and Methods, section 2.12. Burial of Solvent Exposure). This fourth burial bin contains the largest fraction of the amino acid residues (~80% or more) consistently throughout the entire MD trajectory in all proteins (**Supplementary Figure S9**).

The ‘complementarity’ calculation was repeated twice: (i) with all atoms of the protein contributing to the electrostatic complementarity (set 1) and (ii) with salt-bridges being neutralized (set 2). For set 2, salt-bridges were identified *in situ* in each MD snapshot, and, the side-chains involved in the salt-bridges were assigned partial charges of the corresponding uncharged amino acids. This methodology was in accordance with the one used earlier to quantify the relative contribution of salt-bridges to electrostatic complementarity (EC) at protein-protein interfaces [71].

Mild variations were noted in the three P_Em_ measures calculated on main-chain, side-chain and the van der Waal′s surfaces of the whole residue. This indicates that the electrostatic balance is evenly retained at different surface patches throughout the entire residue surface without having any bias towards a particular patch (say, the side-chain dot surface points). For all three P_Em_ measures (P(E_m_^all^), P(E_m_^sc^), P(E_m_^mc^)), a consistent trend (Figure 6, **Supplementary Figure S11, S12**) was observed throughout the MD simulation trajectories where the scores in set 2 were always lesser than or equal to their corresponding scores in set 1. This characteristic trend was found for all four proteins and in no case an exception was observed. The impact was more pronounced in IDPRs rather than IDPs (Figure 6), which is in accordance with the prevalence of persistent salt-bridges in the former. It was also interesting to find out that the time evolved all-atom-P(E_m_) values (set 1) for all four proteins were consistently above the suggested empirical threshold of-1.8 (standardized for globular proteins [41]) throughout the entire MD trajectory. For proteins in general, a P(E_m_) value falling below this threshold corresponds to structures with unbalanced electric fields throughout its van der Waal′s surface representing a state of electrostatic dissonance caused by sub-optimal distributions of atomic partial charges in the molecule. The fact that P(E_m_) for the IDPs/IDPRs were consistently above this empirical threshold throughout the trajectory (Figure 6, **Supplementary Figure S11, S12**) reveals the meticulous balance of electric fields coming from different parts of the chains in their respective time-evolved confomational ensembles. In other words, IDPs/IDPRs seem to maintain their global electrostatic balance with time while continuously fluctuating between different conformations. Although, there was an average of ~5% reduction in the P(Em) values when salt-bridges were neutralized (**Supplementary Figure S13**), noticeably, the reduced values did not drop below the P(Em) threshold of-1.8. This means that even without the contribution of the salt-bridges, IDPs/IDPRs manage to retain their global electrostatic balance. However, the ionic bonds unequivocally contribute to the overall balance. Similar results were also obtained at protein-protein interfaces [71] and within folded globular proteins [17] where the electrostatic complementarity was augmented by the contribution of salt-bridges, though it did not reduce drastically even after neutralizing the salt-bridges.

**Figure 6.**
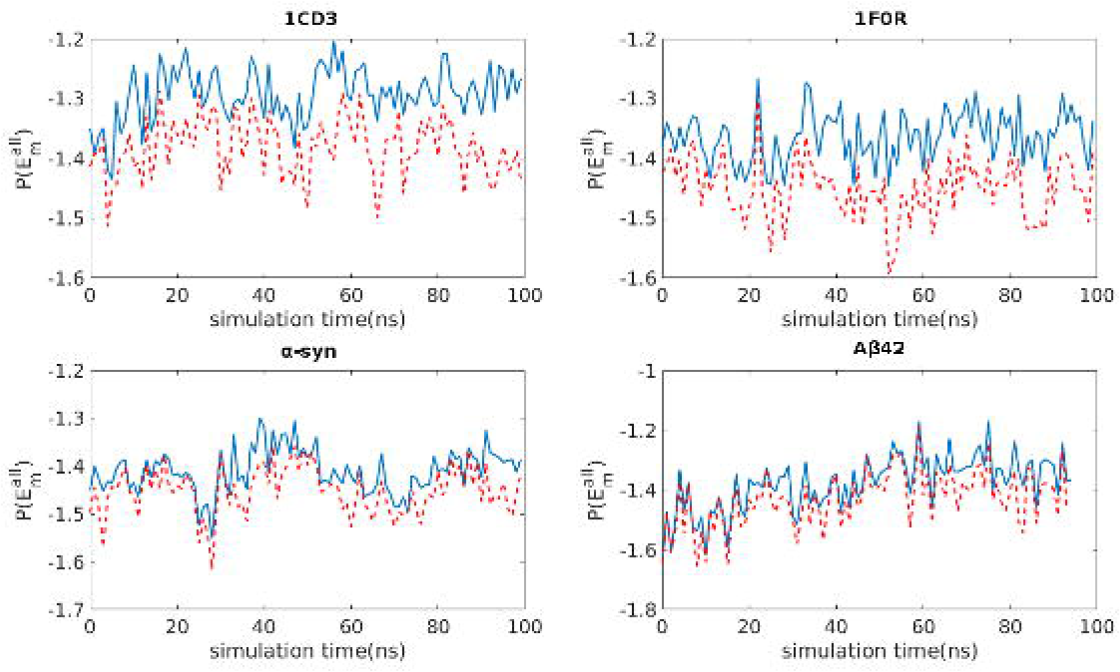
Impact of salt-bridges on the overall electrostatic balance in IDPs as revealed by the time-evolved P(E_m_) score. The log-odd probability score P(E_m_) computed on the distribution of electrostatic complimentarity (E_m_) of the amino acid residues as a function of their burial plotted against simulation time for the four IDPs. This profiles of P(E_m_^all^) are computed on the van der Waal’s surface of the whole-residue. The Prussian blue curves represent the P(E_m_^all^) calculated considering the contribution of all atoms while the orange-red dashed curves represent those calculated when the salt-bridges were neutralized (see **Main Text**).

### 3.7. Correlated movements of oppositely charged residues forming persistent salt-bridges

Dynamic Cross Correlation Maps (see Materials and Methods, section 2.15. Dynamic Cross Correlation Map) were constructed to examine the correlated movements between the oppositely charged residues forming persistent salt-bridges. In consistency with all other calculations, a persistence cut-off of 25% was considered. Traditionally DCCM has been used to portray protein-domain movements between dimers involving whole main-chain trajectories [72,73]. In contrast, our focus was on the dynamic coordination of individual pair of (charged) residues. A majority of the (i,j) pairs recorded high DCC(i,j) values close to 1 (**Supplementary Table S5**) revealing dynamically sustained correlated movements between them. In more detail, 73% and 70% of (i,j) pairs gave rise to a DCC(i,j) value greater than 0.70 in main-and side-chain atoms respectively (Figure 7, **Supplementary Figure S14**). These correlated pairs spanned across ionic bonds of different contact orders, viz., short range contacts (e.g., 3-GLU ~ 5-ARG in Aβ_42_: **Supplementary Table S5**), middle range (e.g., 10-ARG ~ 27-GLU in 1F0R) and long range contacts (2-GLU ~ 108-ARG in 1CD3) with persistence values ranging from 0.257 (viz., 20-GLU ~ 58-LYS in α-syn) to 0.963 (viz., 98-ASP ~ 100-ARG in 1CD3). Only weak correlations were obtained between DCC and persistence (0.33, 0.40 for main-and side-chain DCC respectively) and also between DCC and contact order (−0.40,-0.35). The negative sign (anti-correlation) in the later case indicates that short-range contacts generally have a tendency to show a higher degree of correlated movement-which may be anticipated intuitively. In fact, in 85% (main-chain) and 80% (side-chain) of the cases involving short-range contacts (say, CO ≤ 0.2), a DCC(i,j) value of ≥ 0.70 was observed. However, as pointed out earlier, long and middle range contacts also attributed to high DCC, while, conversely, low / moderate DCC values were also obtained, though rarely for reasonably short-range contacts (e.g., 108-ARG ~ 133-GLU, 82-LYS ~ 97-GLU in 1F0R). It must be noted that 92% of all pairs exceeded a DCC(i,j) value of 0.7 (83% exceeded 0.8) out of the pull of all salt-bridges that have a persistence of 0.5 (i.e., the salt-bridge occurred half the time) or more. Taking into account both the above criteria (i.e., CO ≤ 0.2 and *pers* ≥ 0.5 simultaneously) 100% of all residue pairs forming persistent short-range ionic bonds gave rise to a DCC of 0.70 or more (in both main-and side-chain). Taking all these together, it appears that short-range and persistent ionic bonds generally have a higher tendency to dynamically sustain correlated movements, though, the correlation may not be restricted to any particular sequence separation (i.e., contact order) and/or dynamic spatial separation (i.e., persistence) of the pair of oppositely charged residues. The DCC(i,j) values computed for main-chain and side-chain atomic units (see **Materials and Methods**, section 2.15. **Dynamic Cross Correlation Map**) were also in good agreement, giving a pairwise Pearson′s correlation of 0.97 over thirty persistent salt-bridges (P-value < 0.00001) occurring in all four proteins. This means that the correlated movements generally occur throughout both residues and not restricted to a particular atomic segment. Noticeably, there was only a single instance of weak anti-correlation obtained in the DCC values (77-ARG ~ 105-GLU in 1CD3) continuously throughout both in main-and side-chain atoms.

**Figure 7.**
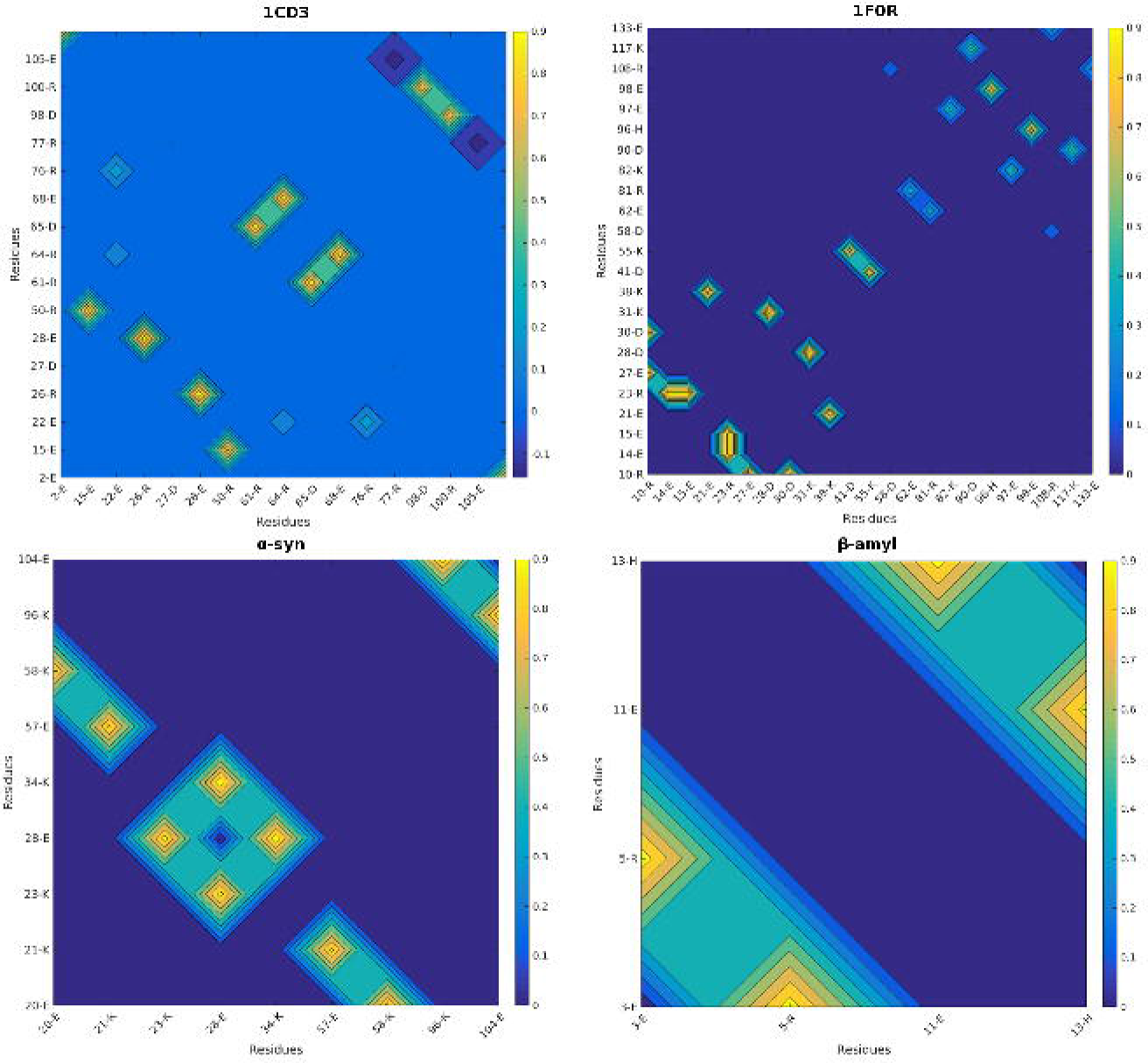
Dynamic Cross Correlation Map (DCCM) composed of the pair of oppositely charged residues forming persistent salt-bridges alone (calculated on their main-chain atoms). Entries apart from the relevant ones (i.e., the salt-bridge forming pair of oppositely charged residues) in the matrices are zero-padded. Amino acid residues are represented (along the horizontal and vertical axes) by their sequence residue numbers (in an ascending order) with their one letter abbreviation codes. Values are plotted according to the color-bars in each sub-plot. Note that only 1CD3 has a negative entry and thus the corresponding color-bar is different than that of the other three.

### 3.8.1 Correlating native sequence with its conformational flexibility: A case study of salt-bridge-mutations in Aß_42_

To address whether the native sequence is indeed the key determinant of the salient dynamic features of the salt-bridges, a case study was performed comprising of short (30 ns) simulations performed on two appropriately chosen multiple point mutants of Aß_42_ (Aß_42_.M1, Aß_42_.M2) (see **Materials and Methods**, section 2.18. **Design of salt-bridge mutants in Aß_42_**). The mutation-approach, implemented to study IDPs, has traditionally been devoted towards exploring their molecular evolution [74] and their pathological features [75]. In fact, there exists a good volume of literature on mutations performed on Aß_42_ itself [76-83], designed to be located on the N-terminus and the middle part of the protein chain. These mutations, mostly ‘hereditary’ [75], were chosen based on geographic and ethnic variations, pedigree of individual families with a history of the Alzheimer′s disease, and probed by techniques directly related to the disease pathology (e.g., fibrillogenesis, aggregation kinetics, neurotoxicity etc.). Even here, salt-bridges were revealed to be influential in mediating the ‘mutation-induced rigidity’ associated with enhanced aggregation of the protein-as recently pointed out in a comprehensive review [75]. In the current case study, the essential idea, however, was to explore the conformational plasticity of IDP′s due to salt-bridge dynamics that may possibly serve to benefit IDP-functions. In other words, the object of the exercise was to investigate whether the salient features of salt-bridge dynamics as characterized in the native conformational ensembles (i.e., the biologically active forms) remains preserved even after the mutations.

Four carefully chosen charged residues were mutated to alanine in both mutants (see **Materials and Methods**, section 2.18. **Design of salt-bridge mutants in Aß_42_**). In one of the mutants (Aß_42_.M1) these mutated residues were the ones that formed high persistence salt-bridges in the native, while, in the other (Aß_42_.M2), the residues involved in multiple transient ionic bonds were mutated. In effect, two high persistence and ten transient salt-bridges were dismantled in the two mutants respectively. In both mutants, no two residues selected for mutation were common. Except for 28-Lys (in Aß_42_.M2), the other seven point mutations were located on either the N-terminus or the middle part of the native Aß_42_ sequence-from where most Aß_42_ mutations are chosen traditionally [75]. In all analyses, an equivalent patch of the simulation trajectory (30 ns from the beginning) of the native was used as a reference.

The analysis of the shape factor profiles in both mutants exhibited lower estimates of ρ fairly consistently throughout their simulation trajectories with respect to the native (**Supplementary Figure S15**) structure. The difference was more prominent in the high-persistence salt-bridge mutant (Aß_42_.M1) as compared to the transient salt-bridge mutant (Aß_42_.M2). The relative shape-factor profiles are indicative of the fact that the mutants populated more compact states relative to the native one. The percentage drop in the ρ values ((max(ρ)-min(ρ))/max(ρ)) also portrayed the same trend, wherein, the fluctuations were the least in Aß_42_.M1 (22.7%), followed by that in Aß_42_.M2 (28.7%) while the native recorded distinctly greater fluctuations than both (35.3%). This trend is perhaps anticipated as the charged moieties are substituted by hydrophobic methyl groups (alanine) in both mutants, leading to reduced electrostatic interactions, potentially promoting the chances of hydrophobic assemblage. In effect, the conformational flexibility was also compromised in the mutants-which were tested by two further calculations, to be discussed next.

#### 3.8.1.1. Sliding of a high-persistence salt-bridge along the sequence due to mutation

First, salt-bridges were identified in both mutants along their respective simulation trajectories. The persistence of these salt-bridges were calculated and compared with the native salt-bridge profile corresponding to the same time interval. In the high-persistence mutant (Aß_42_.M1), only one high-persistence salt-bridge was encountered (**Supplementary Table S6**), that comprised of two adjacent residues, 6-Hip and 7-Asp *(pers*: 0.415). This salt-bridge was also present in native, albeit with somewhat lower persistence *(pers*: 0.183) which is understandable as the two residues were surrounded by several other charged residues to promote possible alternative ionic bonds. Four transient salt-bridges were also found in Aß_42_.M1, but, notably, with a lesser degree of involvement in multiple pairs.

Aß_42_.M2, on the other hand, presented a very interesting case with the sliding of a high persistence salt-bridge along the sequence. In other words, instead of the native high persistence salt-bridge, 3-Glu~5-Arg *(pers*: 0.878), the salt-bridge that was stably formed in this mutant was between 5-Arg~7-Asp with a sliding of two-amino-acids along the sequence. Note, that 3-Glu was not among the mutated residues in Aß_42_.M2. The other persistent salt-bridge in the native, namely, 11-Glu~13-Hip *(pers*: 0.266), however, remained intact with a similar persistence *(pers:* 0.335). A closer look into the ensemble of structures revealed the precise role of the native sequence for this sliding. It is to be noted that two of the four mutations in Aß_42_.M2 (1-Asp, 6-His: see **Materials and Methods**,section 2.18. **Design of salt-bridge mutants in Aß_42_**) were close in the sequence space and towards the N-terminus where all the high-persistence ‘short-ranged’ salt-bridges, both in the native as well as in the mutants, were found to occur. Indeed, these two mutations have a major impact on the salt-bridge sliding. The electrostatic repulsion on 3-Glu due to its neighboring like-charged residue, 1-Asp seemed to enforce its conformational selection, best suited for the stable formation of 3-Glu~5-Arg. The structural context of this ionic bond then influences the formation of 6-Hip~7-Asp (native). Now, in the mutant, Aß_42_.M2, 1-Asp being mutated to alanine, completely eliminates the scope of the aforementioned electrostatic repulsion on 3-Glu originating from the N-terminus. Again, the lack of a bulky and/or charged side-chain in the 6^th^ position (6-Hip → 6-Ala) also facilitates the conformational selection for both 5-Arg and 7-Asp to form a stable ionic bond. Notably, 3-Glu, faces outward opposite to the 5-Arg conformer (Figure 8) in the mutant, with a greater degree of freedom.

**Figure 8.**
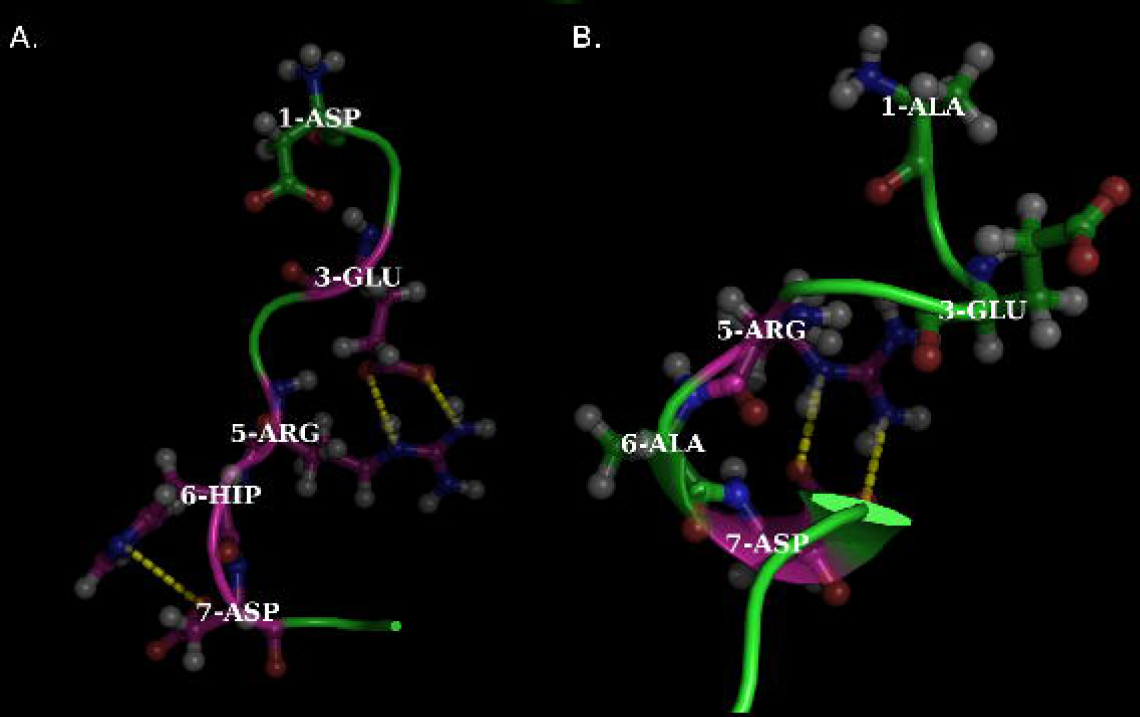
Sliding of a high-persistence salt-bridge along the sequence due to mutations. Panels (A) and (B) represent the native Aß_42_ and the mutant Aß_42_.M2. Note that the high persistence salt-bridge 3-Glu~5-Arg (native) is absent in the mutant and is replaced by 5-Arg~7-Asp which was not present in the native. Note the orientation of 3-Glu is opposite to that of 5-Arg in the mutant (See Main Text for explanation) and that 7-Asp is involved in another salt-bridge (with 6-Hip) in the native.

So, indeed it appears that the ‘native’ sequence is the key determinant of the precise nature and type of salt-bridges (i.e., whether, persistent or transient) with an associated conformational plasticity offered by the particular (native) set of ionic bonds. The process is also naturally coupled with conformational selection of the corresponding elongated charged residues (out of all possible pairwise rotameric combinations), involved in high-persistence ionic bonds.

#### 3.8.1.2. Accounting for the conformational variations in the mutants compared to the native ensemble

In order to probe directly whether the mutations were indeed accountable for a possible reduction in the protein conformational space (as revealed from their dynamics), the requirement was that of a metric to quantify the variation in the coordinates upon structural superposition of the frames. Traditionally the simplest measure is of-course the root mean square deviations (rmsd) in backbone-(or C^a^-) atomic coordinates. To that end, the 30 ns simulation trajectories of the respective mutants and the corresponding interval of the native structure were first split into 6 epochs of 5 ns (0-5, 5-10 and so on…). The two middle epochs (10-15 ns, 15-20 ns) were chosen as templates for superposition and the rmsd’s were recorded. In all three sets (the native and the two mutants), a total of 600 frames were sampled at an interval of 25 ps, covering the whole 30 ns trajectory, and superposed, in-turn, on each of the 200 equidistant snapshots (templates) extracted from the middle epochs (i.e., 10-15, 15-20 ns, sampled at an identical interval). Identical pairs of templates and models were ignored. This produced a rmsd matrix of dimension (((600×200)/2)-200) = 59800, certainly large enough to do meaningful statistics. Results were consistent for both backbone-and C^a^-rmsds.

It was clear and unambiguous from the relative distributions of the C^a^-rmsds that the native structure displayed greater variations in its conformational ensemble than both mutants (Figure 9). As is reflected in Figure 9, the native distribution was multi-modal with three distinctly different peaks, with the major mode at 76.68 Å with another higher-valued mode at 100.42 Å. Only the first native-mode, mapping to conformations with appreciably less variations (mode-value: 9.38 Å) overlapped with the distributions obtained from both the mutants. On the other hand, the high-persistence-salt-bridge-mutant (Aß_42_.M1) produced a unimodal and approximately symmetric distribution (mode: 11.69 Å, mean: 13.13 Å, median: 13.03 Å) with the least conformational deviations among all the three sets. While the transient-salt-bridge-mutant (Aß_42_.M2) had one overlapping mode (at 8.12 Å) as that of Aß_42_.M1, corresponding to comparatively low-variant conformers, it also had another excrescent higher-valued mode (at 26.44 Å), that represented comparatively greater conformational variations.

**Figure 9.**
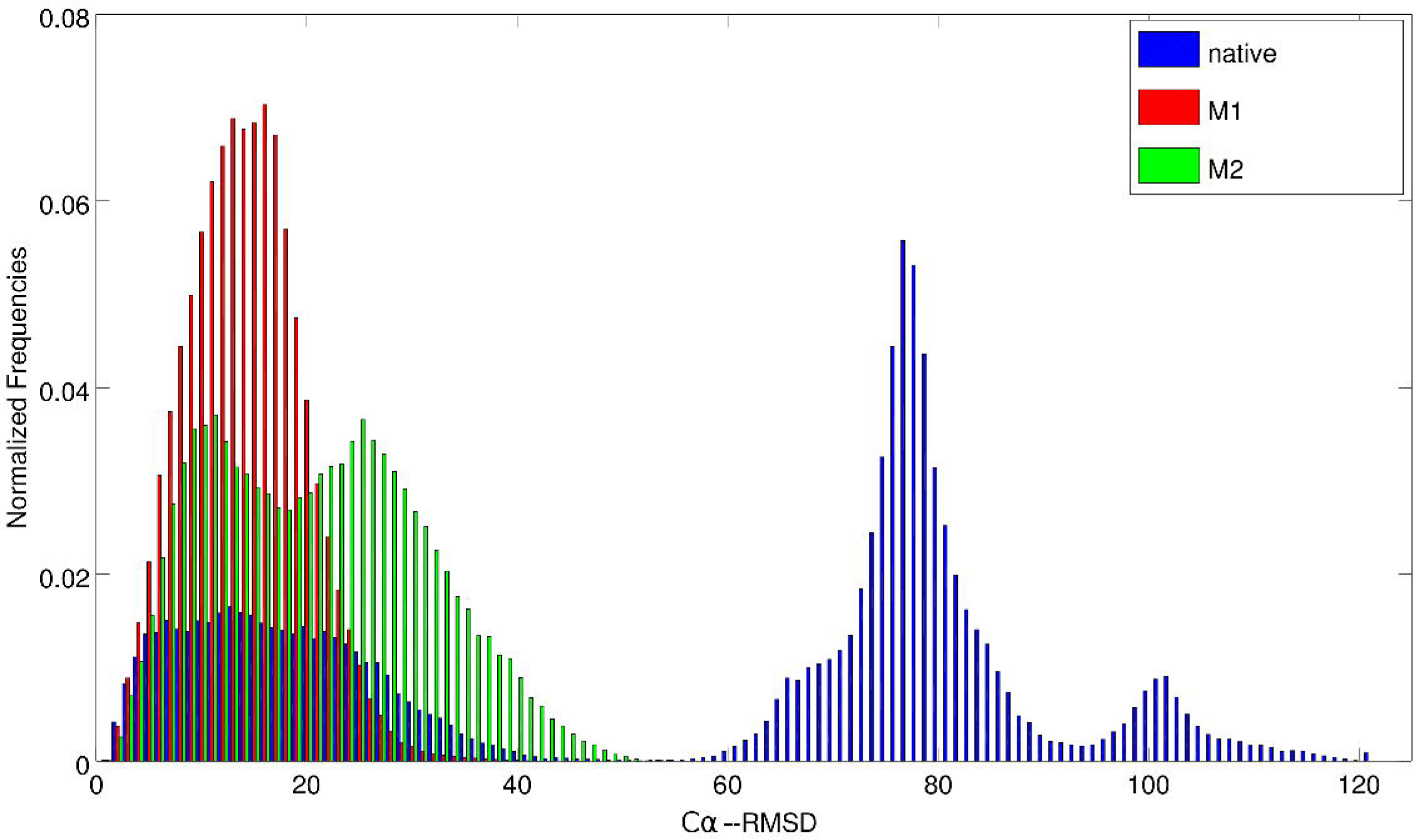
Normalized frequency distribution of C^α^-rmsd values obtained from the native (AB) and the mutant ensembles, Aß_42_.M1, Aß_42_.M2. Note that the native distribution is distinctly different and multi-modal as compared to the unimodal distributions of the mutants (see Main Text for details).

For cross-validation, the analysis was further complemented by the template modeling score (TM-score) [84], a rotation matrix based measure, part of the popular sequence-independent-structural-alignment tool, TM-align [85]. TM-score is based on the principles of inverse square scaling operated on distances between aligned residues, and, in-effect, is an ascending function (i.e., higher the better) for assessing structural similarity between two given atomic models, in contrast to the descending nature of rmsd (lower the better). In contrast to the unbound nature of rmsds, TM-score is defined in the range [0,1], and, returns a trivial value of 1 for two identical protein chains. While a threshold of 0.5 [85] is used to cluster models defined to be in the same fold, a TM-score value of 0.17 is obtained for an average pair of randomly related structures [85]. Overall, higher the average TM-score, better is the similarity between atomic models in an ensemble, and, on the other hand, lower average TM-scores would represent greater degree of conformational variations in an ensemble.

The average TM-scores (in a population of 59800 matrix entries) for both mutants (Aß_42_.M1: 0.340 (±0.108); Aß_42_.M2: 0.283 (±0.068)) were found to be appreciably higher than the corresponding native values (0.260 ±0.060), that amounts to an increase of ~11-18% when normalized by the native standard deviations. In contrast to the distribution of native rmsds, all TM-score distributions (for both mutants and the native) were found to be unimodal, allowing us to perform a t-test. A two tailed p-value < 0.0001 for both the pairwise distributions (Aß_42_.M1 vs. Aß_42_; Aß_42_.M2 vs. Aß_42_) confirmed that both the differences were significant at 0.01% level (i.e., 99.9% confidence) of significance. The statistics clearly implies that there is definite reduction in the conformational variation corresponding to the mutants compared to the native ensemble. The results from both measures, rmsds and TM-score, are in good accordance with each-other.

### 3.8.2 Plausible impact of salt-bridge dynamics in IDP-functions: A docking case study

Having characterized the essential dynamics of salt-bridges in IDPs, it was also important to conceive whether these salient features may be envisaged as (at least partially) being ‘optimized’ for their function. To that end, we selected α-synuclein, an IDP with a wide variety of known functions and malfunctions and performed a systematic case study of docking with one of its well known partners, Tubulin. The misfolded form of α-synuclein forms insoluble fibrillary aggregates, also known as the ‘Lewy bodies’ leading to the Parkinson’s disease [86]. The protein, however, in its monomeric functional form binds to tubulin, arguably its most well studied binding partners [47], influencing the polymerization and dynamics of microtubules, essential for intracellular transport, metabolism, and cell division.

The tubulin-α-synuclein binding has been well-characterized by *in-vitro* pull down assays, co-immunopreciptation and confocal microscopic studies with the full-length as well as the GST-fused truncated domain constructs of α-synuclein. In particular, it has been categorically shown by confocal microscopic co-localization assays that the full-length α-synuclein interacts with tubulin, while the primary interacting region has been mapped to the core (NAC) domain (residues 60-95) [47]. The NAC domain has a higher hydrophobic content (48.6%) than both the overlapped N-terminal (residues: 1-87: 41.4%) and the C-terminal (residues: 96-140: 26.5%) domains. Also important to note that a fairly long patch in the NAC domain (from residues 61 to 79) is devoid of any charged residue(s). In spite of mapping the tubulin-interaction site in α-synuclein, not much is known (to the best of our knowledge) about the binding mechanism from a structural viewpoint. Again, the N-terminal domain of the protein consists of amphipathic repeats with Lysine-rich basic regions (net-charge: +3) and the C-terminus is acidic (net-charge:-12), giving them a fair chance to come in close proximity by virtue of the characteristic dynamic bending due to salt-bridge dynamics and other long-range electrostatic interactions, as has been revealed from the shape factor analysis (section 3.4. **Dynamic bending of IDP chains**).

Thus, one of the major points of interest in this docking study was to explore whether this dynamic bending was influential in the docking (or anchoring) of the middle NAC domain onto tubulin. Docking was performed (in Autodock4 [48]) with the full-length α-synuclein (which is the biologically active form of the protein) as receptor and tubulin as ligand (see **Materials and Methods**, section 2.17. **Molecular Docking**). The choice of α-synuclein as the receptor was based on the fact that it had some prior binding information (that the NAC domain maps to the interaction site) [47] while tubulin had none. All the docking runs were operated in their full *ab-initio* mode (i.e., blind docking), wherein, as a default, the center of the grid-box was superposed to the centroid of the receptor. Since this centroid falls onto the NAC domain (for its middle location in the sequence), the docking initiation site was thus automatically mapped to the NAC domain, even in the default protocol of the blind docking.

As also illustrated in **Materials and Methods** (section 2.17. **Molecular Docking**), the whole 100 ns trajectory of α-synuclein was first equally divided into 5 epochs of length 20 ns each (i.e., 0-20 ns, 21-40 and so on…) and 20 equidistant frames (at an interval of 1 ns) were sampled in each epoch to serve as receptor-templates in 20 independent docking runs. Hence, each epoch was represented by an ensemble of conformations, effectively sampling the whole period. Thus, 100 independent docking runs were carried out equally divided into 5 epochs, wherein, each run again consisted of 100 cycles of the docking algorithm (Autodock). The lowest energy conformers among the 100 cycles were then selected for further analyses, wherein, inter-chain salt-bridges between the two protein chains (α-synuclein and tubulin) were identified in each selected docked pose. All statistics were initially carried out individually in each 20ns epoch with no overlap between adjacent epochs and all inter-chain salt-bridges were considered which occurred at least once in an epoch. For the salt-bridge analyses, the variations in the relative orientations of the selected docked poses were ignored, and in staid, emphasis was given on whether any general recurrent pattern in the salt-bridge dynamics were evident in the whole ensemble, composed of (twenty) docked poses in an epoch. To that end, further analyses involved the identification of the charged residue partner in the inter-chain interaction of the salt-bridges of α-synuclein and recalling their ionic-bond-status (i.e., whether the they are involved in persistent or transient salt-bridges) in the free-form of the protein. It was indeed interesting to find that the majority of these inter-chain salt-bridge residues of α-synuclein were involved in multiple salt-bridges with a wide variety of persistence in the free¬form of the protein. A combined statistics of this multiplicity (i.e., the number of different alike-charged-pairs involved in the free form) in the whole 100 ns trajectory returns an unimodal, approximately bell-shaped distribution giving the peak value at four alike-charged-pairs with a corresponding fractional count of 43.8% (**Supplementary Figure S16F**). Inter-chain-ionic-bond residues involved in single (high-persistence) salt-bridges in the free form of the protein were marginal (3.13% of the whole accumulated population; 0, 8.7, 0, 4.2 and 0% respectively in the five epochs in their ascending order of simulation time).

But for the first epoch (0-20 ns) similar distributions were obtained in all other epochs (**Supplementary Figure S16B-E**). Understandably, the first epoch (0-20 ns) stood out somewhat different than the others, being the initial phase of the post-equilibrium molecular dynamics, with the largest conformational deviations (as reflected in the corresponding shape factor profile; section 3.4. Dynamic bending in IDP chains). Visual investigation revealed an open conformer of α-synuclein in this epoch (Figure 10A) with the disordered NAC domain pervading through the helices of tubulin. Salt-bridges in this epoch, found on either side of the NAC domain, were largely intra-chain, that remains unperturbed in the free form of the protein. They were generally of high persistence (96-Lys ~ 104-Glu: 0.937; 21-Lys ~ 57-Glu: 0.509; 23-Lys ~ 28-Glu: 0.398; 28-Glu ~ 34-Lys: 0.379), more prevalent on the N-terminal domain, which has amphipathic repeats. Understandably all of them consisted of short-range contacts (Figure 10A). However, few inter¬chain salt-bridges were formed between tubulin and α-synuclein too, during the later half of the epoch (infrequently, after ~10 ns), and these were the ones that mapped to four or more (multiple) charged-pairs in the free form of the protein (**Supplementary Figure S16A**). Note that the middle region of NAC (residue 61-79) is totally devoid of any charged residues.

**Figure 10.**
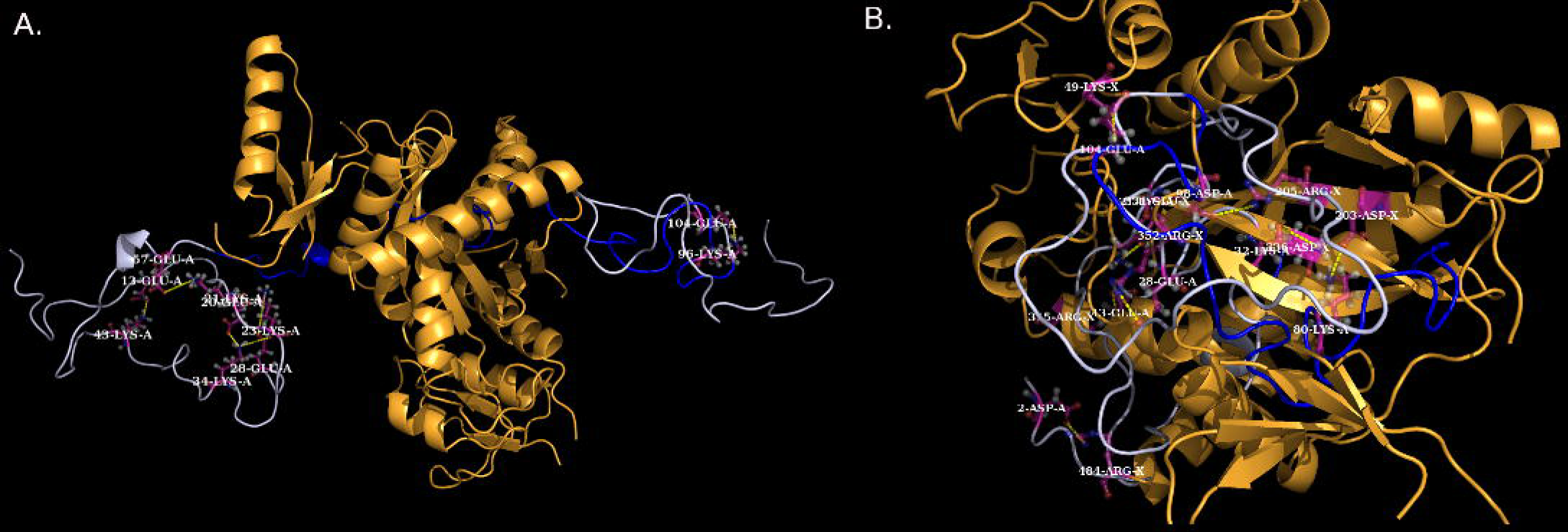
The (A) open and (B) closed conformers of α-synuclein (chain A) docked onto Tubulin (chain X) (drawn in golden yellow ribbon). The NAC domain of α-synuclein is highlighted in blue while the rest of the disordered chain is in light blue. The open form is sampled from the 0-20 ns epoch while the closed docked pose is sampled from the 60-80 ns epoch. Salt-bridges are displayed in dashed yellow line while the involved charged residues are in sticks (colored atom wise: cpk).

From 20ns onward, the docked ensembles corresponding to each epoch revealed dynamically bent disordered conformations of α-synuclein bound to tubulin in closed forms (Figure 10B). Again, in these closed conformers, the disordered α-synuclein chain appears to pervade through the helices of tubulin, followed by tethering of its N-and C-terminal domains, eventually leading to the intertwining of the two chains. The closed conformation especially in context of binding to globular proteins seems plausible, as past studies [60] have also characterized IDPs to populate compact states due to hydrogen bonds and salt bridges rather than to remain in the preferred extended forms.

The desired dynamic flexibility required for this ‘open-to-closed’ conformational conversion (Figure 10) indeed seem to support the salient features of the characteristic salt-bridge dynamics in IDPs. In other words, both wide range of persistence and the multiplicity exhibited in the choice of ionic-bond pairs in the salt-bridges appear to be beneficial for the conformational shift. Furthermore, the restructuring of the salt-bridges from intra-to inter-chain ionic bonds appears to have a more direct impact on the closure of the docked poses (Figure 10B) around the docking initiation site (i.e., the NAC domain). Majority (96.9%) of these salt-bridges were formed by residues that form ‘transient’ ionic bonds in the free protein (α-synuclein), which comprises of multiple charged ionic pairs at different snapshots of the simulation trajectory. The ionic bond traits in the docked poses appear to be characteristic and recurrent (throughout all epochs of 20-100 ns) irrespective of their variations in relative orientations. Thus, indeed, it appears plausible that the characteristic salt-bridge dynamics has both a subtle and a direct influence on IDP-functions, wherein, the interchangeable promiscuous nature of the transient salt-bridges makes them amenable to the exchange of their ionic bond partners in the vicinity of their potential binding partners, thereby promoting inter-chain salt-bridges.

## 4. Conclusions

The main objective of the current study was to understand the basis of a meticulous trade-off between accommodating oppositely charged residues and retaining the desired dynamic flexibility in IDPs/IDPRs. Accordingly, the IDP sequences were chosen such that there are enough oppositely charged residues. The results of this study confirm that the key to maintain such a trade-off lies in the continuous time evolution of salt-bridges with a wide range of persistence. The salt-bridges in IDPs/IDPRs span across the full scale (0 to 1) of persistence values, ranging from extremely short-lived ionic bonds to those that persist consistently throughout the entire simulation trajectory. Moreover, for the transient ionic bonds, a high degree of promiscuity is exhibited by charged groups (say,-NH3+), in terms of pairing to different oppositely charged groups (say,-COO-) originating from a variety of amino-acid residues. Study of the contact order in salt-bridges also revealed the occurrence of short-range, middle-range as well as long-range contacts with no distinct correlation with their corresponding persistence values. In other words, the salt-bridges composed of either closely spaced residues or residues that are distant in the sequence space may correspond to high persistence (say, *pers ≥* 0.8). All these parameters collectively portray a model of dynamic flexibility which was also confirmed visually. Such a model can accommodate oppositely charged residues by allowing the formation of salt-bridges with varying degree of persistence and different ranges of contact orders, and, thereby, map structurally to an ensemble of distant conformations rather than minimally fluctuating about a single global-minima structure as in globular proteins. The ionic bond interactions also appear to be pivotal for the dynamic bending in the IDPs/IDPRs and contribute substantially to their global electrostatic balance. Furthermore, the analyses of hydrophobic burial profiles present a simple and effective single-parameter-tool, the *rGb* score to characterize the potential instability of IDPs/IDPRs in solution which may account for their high reactivity. Interestingly, the charged residues involved in persistent salt-bridges appear to move in a correlated manner throughout the entire dynamic trajectory. This trend was more prominent for short-range contacts with higher persistence. In addition, the study also identified and characterized the dynamics of the composite salt-bridge motifs in the conformational ensembles of the IDPs/IDPRs which mapped the secondary structural transitions to the salt-bridge dynamics. The case studies performed on salt-bridge mutations and docking also suggest plausible ways by which salt-bridge dynamics may influence IDP-functions. This work thus constitutes an important knowledge-base in protein design. So to speak, the results and conclusions of the current study should provoke future studies to explore the disordered-globular interface in proteins and their evolutionary relationship based on salt-bridge design, leading to a mutually reversible switch.

## Acknowledgment and Funding

The work was supported by the Department of Science and Technology-Science and Engineering Research Board (DST-SERB research grant PDF/2015/001079/LS). SB thanks Dr. Pooja Rani and Mrs. Leena Aggarwal for sharing the simulation data of IDP′s.

## Footnotes

1 3.8 Å is the carbon-carbon van der Waals distance

2 A network where every node is connected to every other node

3 μ: mean, σ: standard deviation

